# TGF-β signaling and Creb5 cooperatively regulate Fgf18 to control pharyngeal muscle development

**DOI:** 10.1101/2022.05.25.493396

**Authors:** Jifan Feng, Xia Han, Yuan Yuan, Courtney Kyeong Cho, Eva Janečková, Tingwei Guo, Siddhika Pareek, Jing Bi, Junjun Jing, Mingyi Zhang, Thach-Vu Ho, Yang Chai

## Abstract

The communication between myogenic cells and their surrounding connective tissues is indispensable for muscle morphogenesis. During late embryonic development in mice, myogenic progenitors migrate to discrete sites to form individual muscles. The detailed mechanism of this process remains unclear. Using levator veli palatini (LVP) development as a model, we systematically investigated how a distinct connective tissue subpopulation, perimysial fibroblasts, communcates with myogenic cells to regulate mouse pharyngeal myogenesis. Using single-cell RNAseq data analysis, we identified that TGF-β signaling is a key regulator for the perimysial fibroblasts. Loss of TGF-β signaling led to defects in perimysial fibroblasts and subsequently muscle formation in *Osr2-Cre;Alk5^fl/fl^* mice. In particular, a perimysial fibroblast-specific regulator, Creb5, interacts with TGF-β signaling to enable specific activation of perimysial fibroblast-derived signals such as Fgf18. Moreover, Fgf18 supports pharyngeal muscle development *in vivo* and its exogenous expression can partially rescue myogenic cell numbers in *Osr2-Cre;Alk5^fl/fl^* samples, illustrating that TGF-β-regulated Fgf18 signaling is required for LVP development. Collectively, our findings revealed the mechanism by which TGF-β signaling achieves its specificity in defining the perimysial-to-myogenic signals for pharyngeal myogenesis.

## Introduction

Inductive tissue-tissue interaction constitutes a key mechanism for organ development and tissue homeostasis (Souilhol *et al*., 2016; Takahashi *et al*., 2013; Vainio and Müller, 1997). The interaction and communication of myogenic cells with the surrounding connective tissues are also indispensable for muscle morphogenesis. Although the detailed cellular and molecular aspects of this dialogue remain to be elucidated, the connective tissues have been shown to serve as a signal source and ‘pre-pattern’ for muscle differentiation and patterning during development (Michailovici *et al*., 2015; Nassari *et al*., 2017).

At the onset of pharyngeal muscle development, myogenic progenitors migrate from cranial paraxial mesoderm into the center of each pharyngeal arch to form the primary myogenic sites surrounded by cranial neural crest (CNC)-derived cells. Later, in coordination with the development of other CNC-derived craniofacial connective tissues, myogenic progenitor cells in these mesodermal cores migrate to the final myogenic sites while being segregated and patterned into discrete muscle masses anatomically matching muscles in adults (Noden and Francis-West, 2006; Sambasivan et al., 2011; Shih et al., 2008; Ziermann et al., 2018). A distinct regulatory network upstream of myogenesis has been identified during early-stage pharyngeal muscle development (Buckingham and Rigby, 2014; Sambasivan *et al*., 2011), but the regulatory mechanism that establishes the fine-tuned craniofacial muscle anlagen in their final myogenic sites remains unknown (Noden and Francis-West, 2006).

In the final stage of pharyngeal muscle formation, the first and second pharyngeal arch mesodermal cores respectively contribute to the muscles responsible for mastication and facial expression; the muscles in the oropharyngeal region crucial for breathing, swallowing, and speaking are derived predominantly from the fourth pharyngeal arch, including all the pharyngeal constrictors and soft palatal muscles except the tensor veli palatini (Frisdal and Trainor, 2014; Li *et al*., 2019; Michailovici *et al*., 2015; Shiba and Chhetri, 2019; Sugii *et al*., 2017). Notably, interactions between CNC-derived and myogenic cells become more prominent as the CNC-derived cells start to contribute to connective tissues while individual muscles segregate and proceed to their final locations (Ziermann *et al*., 2018).

During this late pharyngeal muscle development in mice, most pharyngeal muscles of the head have already segregated and reached their ultimate locations between E11 and E13.5 (Noden and Francis-West, 2006). In contrast, the soft palatal muscle anlage development starts later, concurrent with the soft palatal shelf primordium formation (Han *et al*., 2021; Li *et al*., 2019). From E13.5 onwards, individual soft palatal muscles, particularly the levator veli palatini (LVP), can be clearly detected segregating from the rest of the pharyngeal muscles and populating the soft palate primordium, patterning and migrating towards the midline in parallel to the lateral-to-medial outgrowth of the connective tissues consisting of the CNC-derived palatal mesenchymal cells (Grimaldi *et al*., 2015; Han *et al*., 2021; Li *et al*., 2019). Therefore, soft palatal muscle development provides an optimal opportunity to study the initiation, segregation, and migration of muscle anlage formation during craniofacial development.

Clinically, the most common congenital craniofacial malformation is cleft lip and/or palate (Vyas *et al*., 2020). When the soft palate is affected, disruption of the soft palatal shelves separating the oral and nasal cavities co-occurs with defects in soft palatal muscle formation and oropharyngeal function (Li *et al*., 2019; Monroy *et al*., 2012). Furthermore, mouse models have demonstrated that loss of key regulators in the perimysial cells, a subset of the palatal mesenchymal connective tissues closely associated with the muscles, ultimately affects soft palatal muscle differentiation and patterning (Han *et al*., 2021; Sugii *et al*., 2017). These findings suggest that CNC-derived perimysial cells possibly contribute to part of the embryonic myogenic niche for craniofacial muscle development at the later stages. Thus, soft palate development can serve as a useful model for studying how perimysial cells regulate late pharyngeal muscle anlage formation through tissue-tissue interactions.

In the present work, we investigated the mechanism that regulates perimysial-to-myogenic communication using the development of the pharyngeal muscle LVP as a model. Using unbiased single-cell RNAseq (scRNAseq) data analysis combined with mouse genetic approaches, we identified TGF-β signaling as a predominant and specific signaling activity that enables the perimysial cells to interact with the adjacent myogenic cells during late pharyngeal muscle development. Furthermore, we identified that a perimysial-specific regulator Creb5 interacts with TGF-β signaling to co-activate the expression of specific perimysial-to-myogenic signals such as Fgf18 to regulate pharyngeal myogenesis. Taken together, our findings reveal that TGF-β signaling and perimysial-specific regulators may cooperatively define perimysial-to-myogenic signals in regulating pharyngeal myogenesis.

## Results

### Unbiased screening using scRNAseq data analysis identifies TGF-β signaling as predominant and specific signaling for the perimysial cells within the soft palate mesenchyme

During late pharyngeal muscle development, myogenic cells of the soft palatal muscles, particularly those of the LVP, can be observed segregating from the existing myogenic site associated with middle pharyngeal constrictor to populate the soft palatal shelves and pattern alongside the connective tissues consisting of the palatal mesenchymal cells (Han *et al*., 2021; Li *et al*., 2019). In parallel, during this stage, our recent scRNAseq analysis identified that the morphologically homologous palatal mesenchyme is also patterned into heterogeneous cellular domains associated with distinct anatomical locations and gene expression profiles (Han *et al*., 2021). We thus hypothesized that the perimysial population, anatomically adjacent to the myogenic cells, serves as a microenvironment or niche for embryonic pharyngeal muscle development. Furthermore, there must be a specific regulatory mechanism that differentiates the perimysial cells from the rest of the palatal mesenchymal cell populations so that the perimysial cells can provide specific signals to define the myogenic site.

Since signaling pathways have been reported to play critical roles in the patterning and fate determination of craniofacial mesenchymal cells (Mishina and Snider, 2014; Neubüser *et al*., 1997; Vincentz *et al*., 2016; Xu *et al*., 2019), we performed an unbiased signaling activity analysis of the cell populations of the soft palate, focusing on identifying key signaling pathways specifically activated in the perimysial cells. Using CellChat’s incoming signaling analysis from scRNASeq data of E13.5-E15.5 soft palate, we identified inferred signaling pathway activities of different cell types based on enriched signaling interactions (Figure 1A-B, Figure 1—figure supplement 1). Notably, the signaling activity of all the palatal mesenchymal cell types was categorized into the same group due to the shared expression of similar signaling pathways (Figure 1B), consistent with their common connective tissue identity. Among this group of pathways, Ephrin-Eph (EphA/EphB) (Benson and Serrano, 2012; Xavier *et al*., 2016), non-canonical Wnt (ncWnt) (Reynolds *et al*., 2019), IGF (Marchant *et al*., 2020), TGF-β (Iwata *et al*., 2011), FGF (Nie *et al*., 2006), and Hedgehog signaling (HH) pathways (Everson *et al*., 2018) were previously identified to be required for the development of craniofacial mesenchymal tissues including palatal mesenchyme, so we reasoned that they are more likely to be functionally required by the perimysial cell fate determination, too. We thus next analyzed the intensities of activities of these pathways in the perimysial cells and found that the TGF-β signaling activity was inferred to be the highest (Figure 1C). Moreover, a known TGF-β signaling downstream target, *Tgfbi* (transforming growth factor-beta-induced protein), was among the top 10 conserved marker genes for the perimysial cells across various developmental stages (Figure 1D), suggesting that TGF-β signaling is not only a predominant signaling pathway but also specifically associated with regulating the perimysial cell fate and maintaining a perimysial-derived signal.

**Figure 1.**
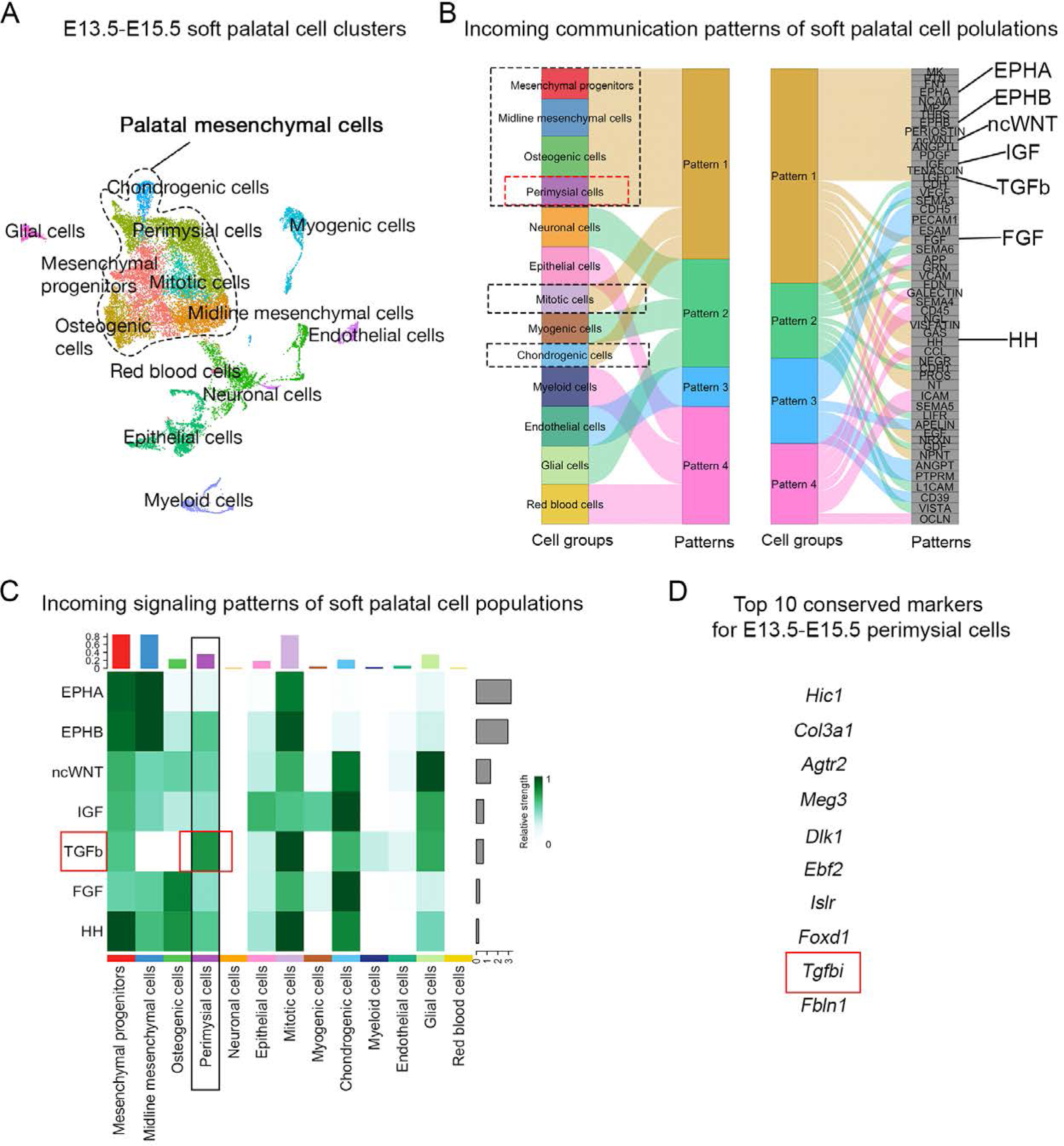
CellChat analysis of E13.5-E15.5 soft palatal scRNAseq data predicts TGF-β signaling as a key pathway activated in the perimysial cells in the developing soft palate. (A) UMAP plot of soft palatal cell populations identified by Seurat integration analysis. (B) Alluvial plot of the incoming signaling patterns for soft palatal cell populations. The black dotted boxed area indicates the palatal mesenchymal cell populations grouped into the same incoming patterns. The red dotted box highlights the perimysial cells. (C) Heatmap of the contribution of the incoming pathways of interest to soft palatal cell populations. The black box highlights signaling activity in the perimysial cells and the red box highlights TGF-β signaling activity in the perimysial cells. (D) Top conserved markers for perimysial cells identified through Seurat integration analysis. This article includes the following figure supplement for figure 1: Figure supplement 1. CellChat analysis of scRNAseq data identified incoming signaling pathways for all cell types in the soft palate.

To validate the signaling activity analysis result from scRNAseq data, we next evaluated the TGF-β signaling activity *in vivo* during different stages of soft palatal development. Consistent with scRNAseq analysis, TGF-β signaling was present in the palatal shelf mesenchyme primordium during soft palatal development from E13.5 onwards (Figure 2A-H) and was most active in the cells surrounding the LVP myogenic cells, anatomically identified as perimysial cells (Figure 2J-L, N-P), particularly during the early segregation and separation process at E13.5 (Figure 2B, F, J, N) and E14.5 (Figure 2C, G, K, O). Notably, TGF-β signaling was ubiquitous in the mesenchymal cells close to the pharyngeal wall at E12.5 prior to the arrival of myogenic cells (Figure 2A, E, I, M). This specific activation of TGF-β signaling in the presence of perimysial cells suggested that TGF-β signaling may play a role in patterning perimysial cells to establish a myogenic niche in the specific anatomical location for the LVP. This postulation is supported by our previous work, which showed that loss of TGF-β in *Wnt1-Cre;Alk5^fl/fl^* mice in the early pre-migratory neural crest cells and their derivatives from E9.5 onward affect proliferation and differentiation of the tongue and other craniofacial muscles (Chai *et al*., 2000; Han *et al*., 2014).

**Figure 2.**
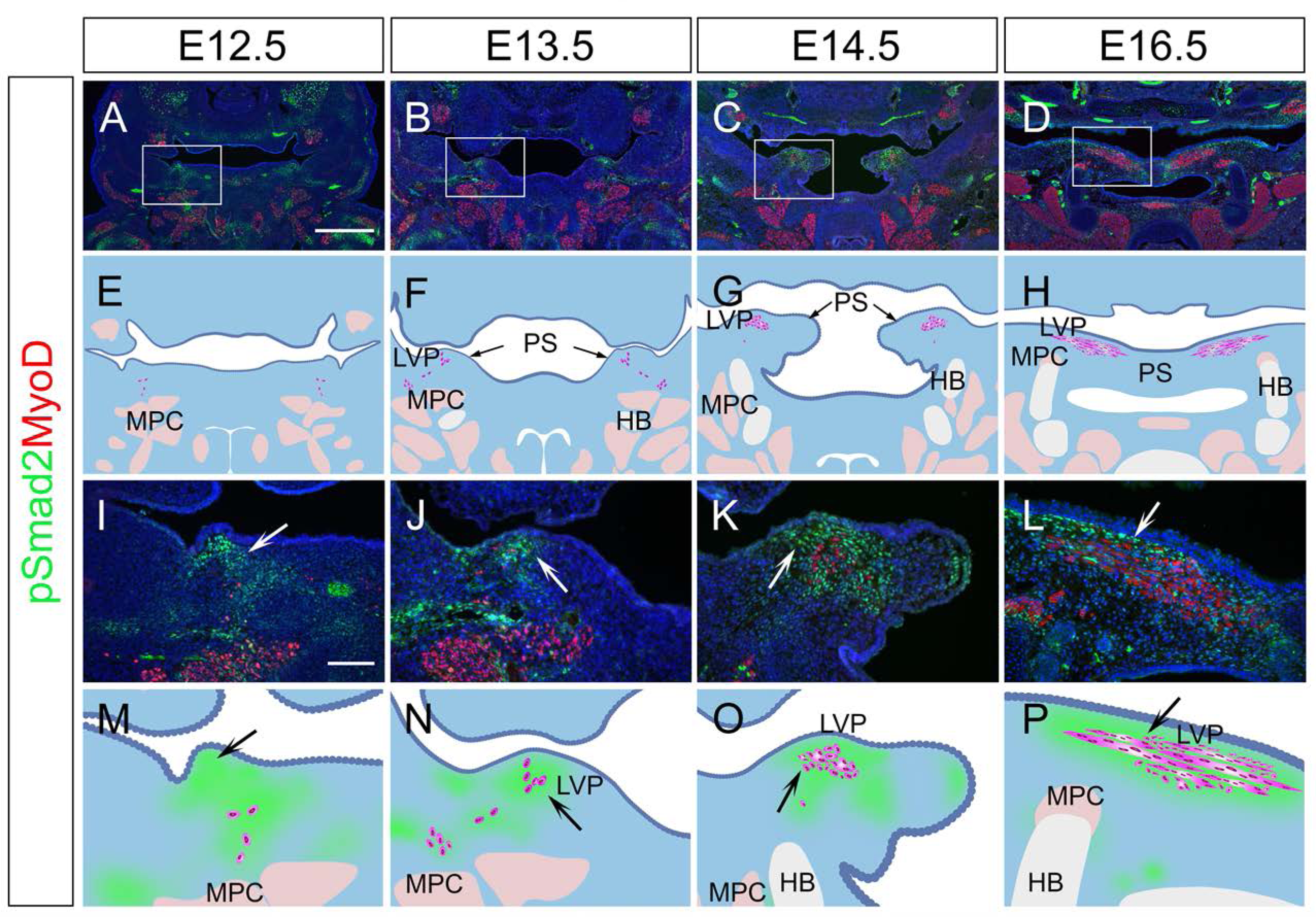
TGF-β signaling is activated in perimysial cells surrounding the developing LVP myogenic cells. (A-P) Immunofluorescence (A-D and I-L) and schematic drawings (E-H and M-P) of myogenic marker MyoD (red) and TGF-β signaling readout pSmad2 (green) expression in coronal sections of LVP region at E12.5, E13.5, E14.5, and E16.5. Boxed areas in A, B, C, and D are shown magnified in I, J, K, and L, respectively. Schematic drawings in E-H and M-P correspond to A-D and I-L, respectively. Arrows in F and G indicate the palatal shelves. White and black arrows in J-L and N-P point to the LVP myogenic sites in the palatal shelves. Arrows indicate the absence of MyoD+ signal in mouse pharyngeal wall at E12.5 in I and M, and the presence from E13.5 onwards in J-L and N-P. The magenta and salmon colors represent the LVP myogenic cells and the other pharyngeal muscles, respectively. HB, hyoid bone; LVP, levator veli palatini; MPC, middle pharyngeal constrictor; PS, palatal shelves. Scale bars in A and I indicate 500 μm in A-D and 100 μm in I-L, respectively.

### **Loss of TGF-β** leads to soft palatal shelf mesenchyme and LVP defects in *Osr2-Cre;Alk5^fl/fl^*

### mice

To test whether this specific TGF-β signaling in the perimysial cells in later developmental stages also plays any functional role in regulating the fate of myogenic cells, we utilized *Osr2- Cre*, which affects the palatal mesenchyme at a later stage than *Wnt1-Cre* (Chen *et al*., 2009). Using *Osr2-Cre;tdTomato* mice, we confirmed that *Osr2-Cre* can specifically label the palatal mesenchyme without affecting the myogenic cells during soft palatal muscle development (Figure 3—figure supplement 1A-D). Thus, we generated *Osr2-Cre;Alk5^fl/fl^* mice, in which TGF-β signaling is ablated in the palatal mesenchyme due to the loss of TGF-β Type 1 receptor *Alk5*. Loss of TGF-β signaling in the palatal mesenchyme, including the perimysial cells, led to morphologically deformed palatal shelves and cleft palate in the LVP region in E18.*5 Osr2- Cre;Alk5^fl/fl^* mice (Figure 3B and D), compared with the intact control palatal shelves (Figure 3A and C), indicating intrinsic defects in the palatal mesenchymal cells. In addition, while abundant MHC+ muscle fibers were present in the control palatal shelves (Figure 3E, I, G, and K), in the defective palatal shelves of *Osr2-Cre;Alk5^fl/fl^* mice, LVP formation was so severely affected that no muscle fibers could be detected at this stage (Figure 3F, J, H, and L). This confirmed that TGF-β signaling is indispensable for the perimysial-derived signaling for myogenesis.

**Figure 3.**
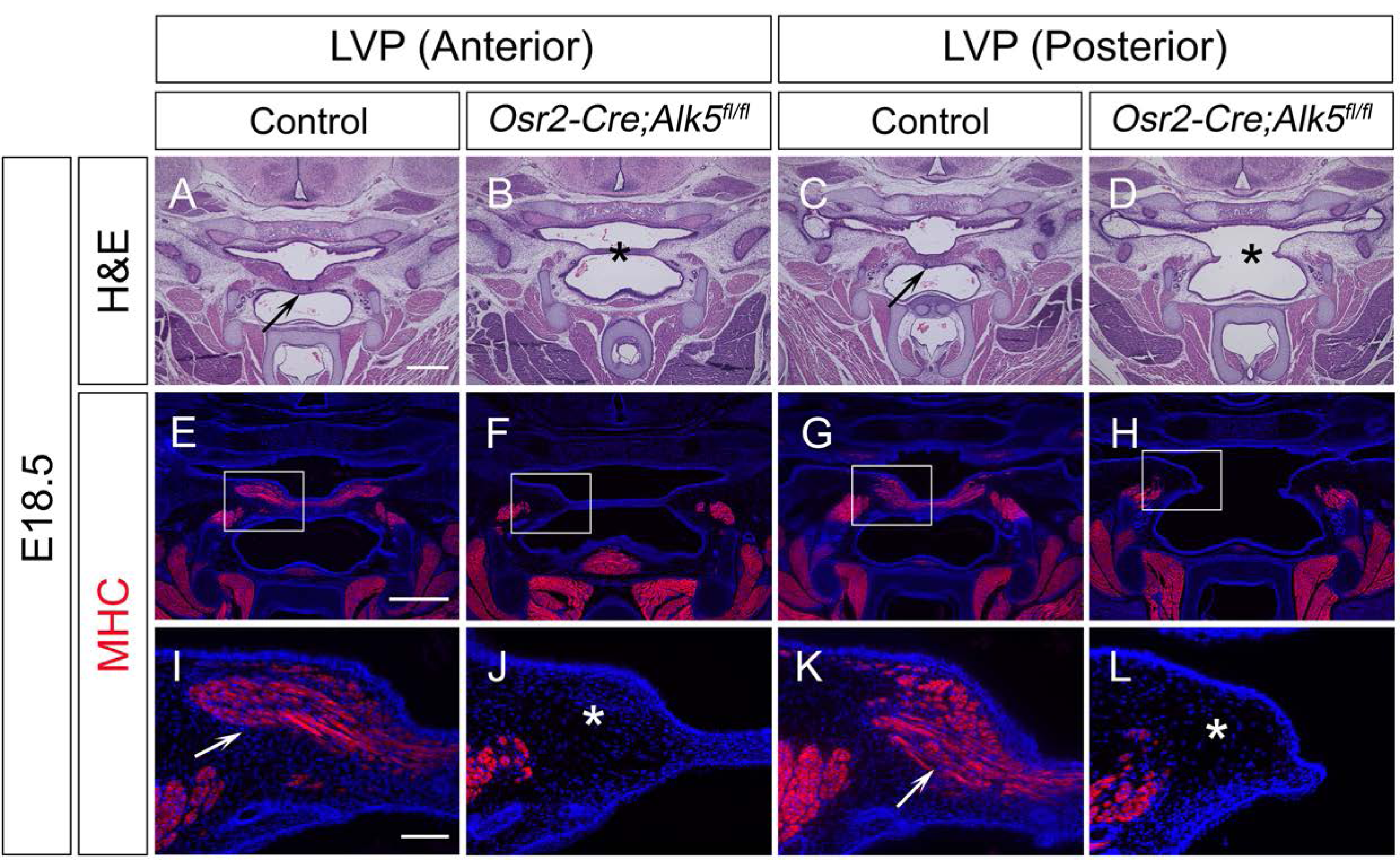
*Osr2-Cre;Alk5^fl/fl^* mice exhibit cleft soft palate with palatal shelf and LVP defects at E18.5. (A-D) H&E staining of coronal sections at LVP region from control and *Osr2-Cre;Alk5^fl/fl^*mice at E18.5. Arrows indicate the normal palatal shelf in A and C, asterisks indicate the deformed and cleft palatal shelf in B and D, respectively. (E-L) Immunofluorescence of MHC (red) in coronal sections at LVP region from control and *Osr2-Cre;Alk5^fl/fl^* mice at E18.5. Boxed areas in E, F, G, and H are enlarged in I, J, K, and L, respectively. Arrows indicate the presence of muscle fibers in I and K; asterisks indicate their absence in J and L. Scale bars in A and E indicate 500 µm in A-D and E-H, respectively; the scale bar in I indicates 100 µm in I-L. This article includes the following figure supplements for figure 3: Figure supplement 1. *Osr2-Cre* targets palatal mesenchymal cells surrounding soft palatal myogenic cells. Figure supplement 2. *Osr2-Cre;Alk5^fl/fl^* mice exhibit palatal shelf and myogenic defects in the LVP region from E14.5 onwards.

We next evaluated the progression of the palate mesenchymal and myogenic cell defects in the *Osr2-Cre;Alk5^fl/fl^* mice. We found that the palatal shelves of *Osr2-Cre;Alk5^fl/fl^* mice started to appear morphologically different from those of the controls at around E14.5, suggesting that defects in the palatal mesenchymal cells start at this stage (Figure 3—figure supplement 2A-H).

In parallel, fewer cells expressed the myogenic marker MyoD in the *Osr2-Cre;Alk5^fl/fl^* mice as early as E14.5 (Figure 3—figure supplement 2A-D). The palate mesenchymal defects became more exacerbated over time, and the myogenic defect also increased in severity over time as myogenic progenitor cells became almost undetectable from around E16.5 (Figure 3—figure supplement 2E-H). Thus, we considered E14.5 as the onset of the defects and used this stage as a starting point for molecular analysis to investigate the mechanism through which TGF-β signaling regulates the perimysial cells. We also confirmed that the TGF-β signaling activity (pSmad2) was efficiently reduced in the perimysial cells adjacent to the LVP in the *Osr2- Cre;Alk5^fl/fl^* mice (Figure 3—figure supplement 2I-L).

### TGF-β signaling is required for *Tbx15+* perimysial fibroblasts in regulating pharyngeal muscle development

To precisely identify the molecular changes of the perimysial population in the *Osr2-Cre;Alk5^fl/fl^* mice, we performed scRNAseq analysis to compare cell-type specific gene expression profiles of cell populations in E14.5 *Osr2-Cre;Alk5^fl/fl^* and control soft palates. Using integration analysis based on shared variance (Figure 4—figure supplement 1A), we identified similar clusters in the soft palates of control and *Osr2-Cre;Alk5^fl/fl^* mice at this stage. We first distinguished the palatal mesenchymal cells (*Runx2+/Twist1+*) from other non-mesenchymal cell types in the soft palate (Figure 4—figure supplement 1B-L). Then, using more detailed markers we recently established from E13.5-E15.5 soft palate scRNAseq analysis (Han *et al*., 2021), we further distinguished the perimysial population (*Tbx15+/Hic1+/Aldh1a2+*) from the remaining palatal mesenchymal populations of the E14.5 *Osr2-Cre;Alk5^fl/fl^* and control mice (Figure 4—figure supplement 1M- N).

**Figure 4.**
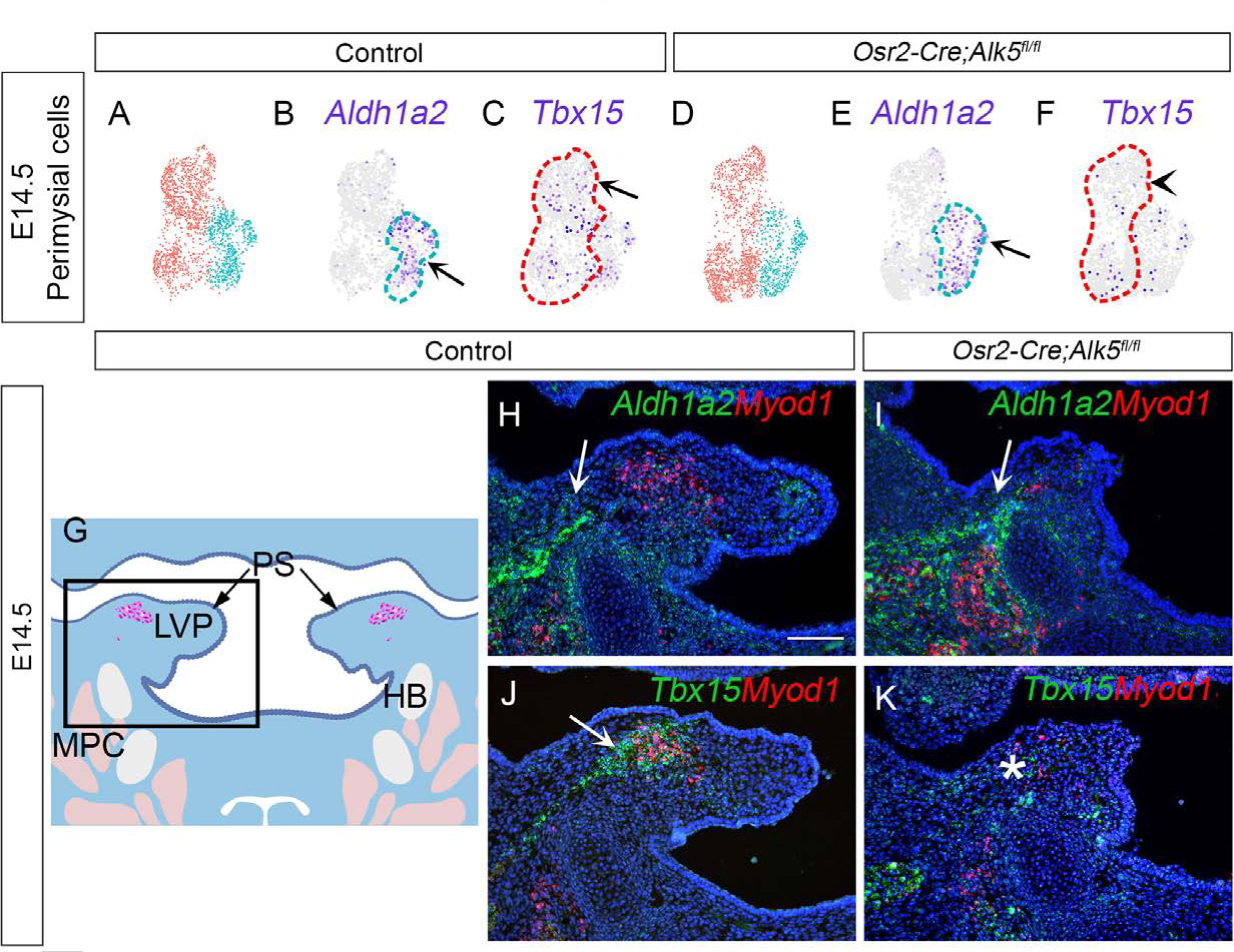
The *Tbx15+* subpopulation is more affected than the *Aldha12*+ subpopulation in *Osr2- Cre;Alk5^fl/fl^* palatal mesenchyme. (A-F) Expression patterns of marker genes *Aldh1a2* and *Tbx15* in early (perimysial progenitors) and late perimysial cell subpopulations (perimysial fibroblasts) at E14.5 in control and *Osr2-Cre;Alk5^fl/fl^*mice. Arrows and arrowhead in B, C, E, and F indicate differential expression of the two genes between control and *Osr2-Cre;Alk5^fl/fl^* mice. Arrows and arrowhead indicate normal and reduced expression level, respectively. (G) Schematic drawing of coronal sections of LVP region at E14.5. Boxed area in G indicates the region of H-K. Magenta and salmon colors represent the LVP myogenic cells and the other pharyngeal muscles, respectively. HB, hyoid bone; LVP, levator veli palatini; MPC, middle pharyngeal constrictor; PS, palatal shelves. (H-K) RNAScope *in situ* hybridization for *Aldh1a2* (green) or *Tbx15* (green) colocalization with myogenic markers *Myod1* (red) in coronal sections of LVP region of control and *Osr2-Cre;Alk5^fl/fl^* mice at E14.5. Arrows in H, I, and J indicate *Aldh1a2* or *Tbx15* expression. Asterisk in K shows reduced *Tbx15* expression. The scale bar in H indicates 100 μm in H-K. This article includes the following figure supplement for figure 4: Figure supplement 1. scRNAseq analysis identified distinct populations of palatal mesenchymal cells in E14.5 *Osr2-Cre;Alk5^fl/fl^* and control soft palates.

Since previous E13.5-E15.5 soft palate scRNAseq analysis inferred that the perimysial cells contain at least two distinct subpopulations with different differentiation statuses, namely earlier perimysial progenitors (*Aldh1a2+*) and the more committed later perimysial population (perimysial fibroblasts) (*Tbx15+*) (Han *et al*., 2021), we sought to identify whether either one or both subpopulations were affected by the loss of TGF-β signaling. By comparing the expression patterns of marker genes for both clusters of the perimysial cells from *Osr2-Cre;Alk5^fl/fl^* and control scRNAseq data at E14.5 (Figure 4A, D), we found the late perimysial cell subpopulation, which accounts for the majority of perimysial cells at this stage, showed a significant reduction of *Tbx15* expression in the *Osr2-Cre;Alk5^fl/fl^* mice (Figure 4C, F), while *Aldh1a2* expression in the perimysial progenitor subpopulation was not affected (Figure 4B, E). Consistent with the scRNAseq data analysis result, *Aldh1a2+* perimysial progenitors and *Tbx15*+ perimysial fibroblasts were found to be associated with different cellular identities *in vivo*. While *Aldh1a2+* mainly labeled the perimysial cells of the adjacent middle pharyngeal constrictor (Figure 4H), *Tbx15*+ cells were located predominantly adjacent to the LVP myogenic cells (Figure 4J), similar to where TGF-β signaling activity has been detected. This overlap between TGF-β signaling activity and the *Tbx15*+ population suggests that TGF-β signaling may be a more specific regulator for the later stage of perimysial cells (perimysial fibroblasts). Consistent with this observation and similar to the scRNAseq analysis, expression of *Tbx15* expression (Figure 4K) was more affected than that of *Aldh1a2* (Figure 4I) in the perimysial cells of *Osr2-Cre;Alk5^fl/fl^*mice. While our recent study showed that Runx2 is a regulator of *Aldh1a2+* early perimysial progenitors (Han *et al*., 2021), here we identified TGF-β signaling as the main regulator for the fate determination of more committed perimysial fibroblasts, suggesting a specific role of TGF-β signaling in late stage-pharyngeal muscle development.

### CellChat analysis of soft palate scRNAseq data identifies putative perimysial-to-myogenic signaling molecules downstream of TGF-β signaling

Since the perimysial fibroblast population is anatomically adjacent to the myogenic cells, they are most likely to be part of a microenvironment that supports myogenesis through signaling communication. To systemically identify potential signaling interactions between these two populations, we examined the E13.5-E15.5 integrated soft palate scRNAseq data with the outgoing signaling analysis of the CellChat package (Han *et al*., 2021; Jin *et al*., 2021) to infer enriched intercellular signaling interactions at the single-cell level. In particular, we focused our analysis on signals sent from perimysial fibroblasts and received by myogenic cells. In doing so, we detected several enriched interactions including those between perimysial-derived signaling molecules in ncWnt, FGF, Notch, and BMP signaling pathways (Wnt5a, Fgf18, Dlk1, and Bmp4) and corresponding receptors expressed in the myogenic cells (Fdz4, Fgfr1/4, Notch3, and Bmpr1a/Acvr2b, respectively) (Figure 5A). We also identified other interactions that may be associated with cell behavior through modulating extra-cellular matrices, such as Thrombospondin (Thbs3-Sdc1), Pleiotrophin (Ptn-Ncl), Nectins (Nectin3-Nectin1), Midkine (Mdk-Ncl), and Laminins (Lama4/b1/c1-Dag1) (Figure 5A).

**Figure 5.**
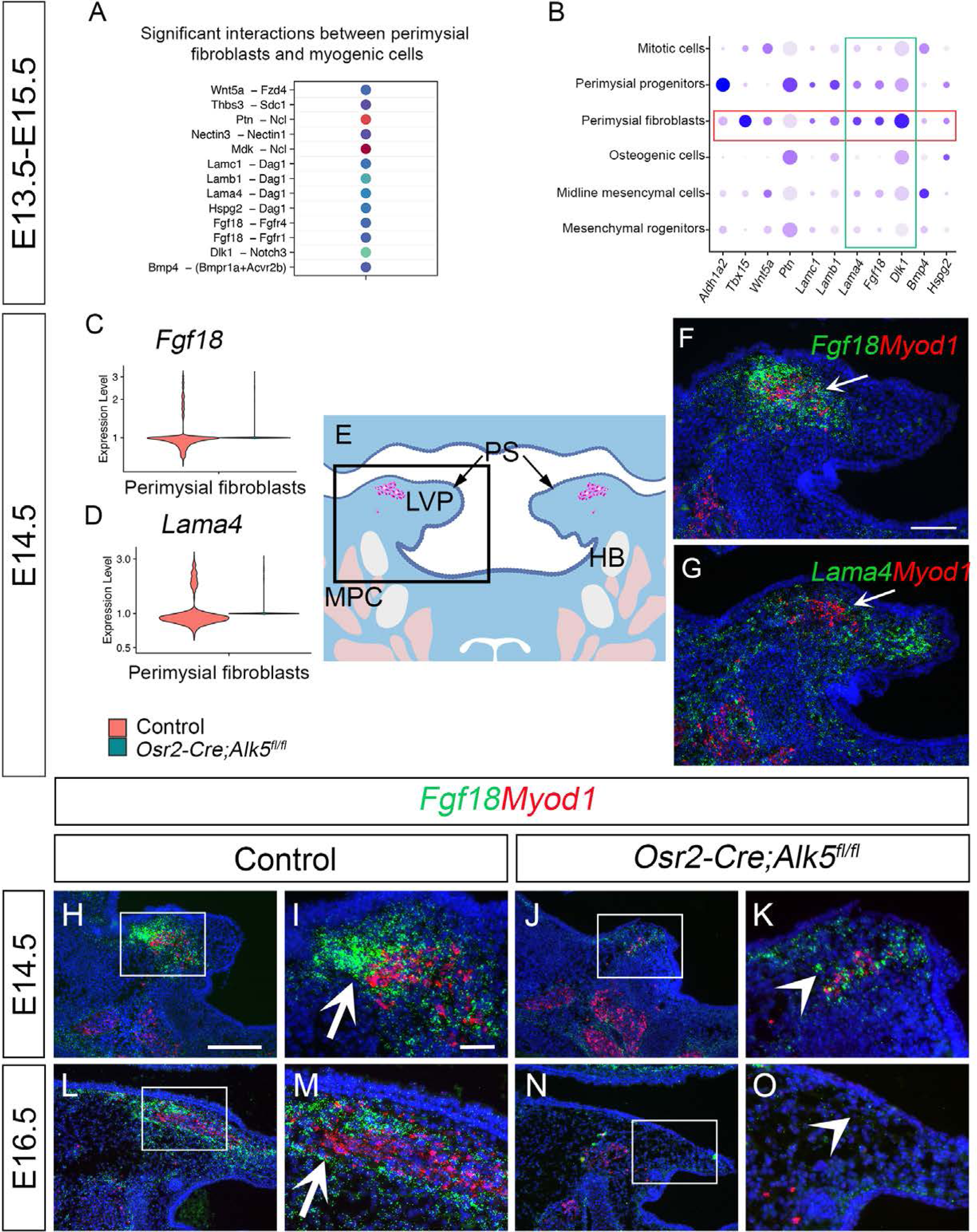
CellChat analysis of soft palate scRNAseq data identifies putative perimysial-to-myogenic signaling molecules also downregulated in *Osr2-Cre;Alk5^fl/fl^* perimysial fibroblasts**. (**A) CellChat analysis of scRNAseq integration data of E13.5-E15.5 soft palatal tissue showing significant interactions between perimysial fibroblasts and myogenic cells. (B) Dot plot of expression patterns of perimysial fibroblast-derived signals in individual cellular clusters of integration scRNAseq from E13.5-E15.5 soft palatal tissue. Note that highlighted signals, *Lama4*, *Fgf18*, and *Dlk1,* are more enriched in the perimysial fibroblasts. (C-D) Expression of *Fgf18* (C) and *Lama4* (D) from scRNAseq analysis of control (red) and *Osr2-Cre;Alk5^fl/fl^* mice (green) at E14.5. (E) Schematic drawing of coronal section of the LVP region at E14.5. Boxed area in E indicates the region shown in F-G. The magenta and salmon colors represent the LVP myogenic cells and the other pharyngeal muscles, respectively. HB, hyoid bone; LVP, levator veli palatini; MPC, middle pharyngeal constrictor; PS, palatal shelves. (F-G) RNAScope *in situ* hybridization for *Myod1* (red) and *Fgf18* (green) (F) or *Lama4* (green) (G) in the coronal section of the LVP region at E14.5. White arrows in F and G point to a positive signal. (H-O) RNAScope *in situ* hybridization for *Myod1* (red) and *Fgf18* (green) in the coronal section of the LVP region at E14.5 (H-K) and E16.5 (L-O) in control (H-I, L-M) and *Osr2-Cre;Alk5^fl/fl^* mice (J-K, N-O). Arrows in I and M point to positive signal; arrowheads in K and O indicate lack of signal. Boxed areas in H, J, L, and N are enlarged in I, K, M, and O, respectively. Scale bar in H indicates 250 μm in H, J, L, and N; the scale bar in I indicates 50 μm in I, K, M, and O. This article includes the following figure supplement for figure 5:Figure supplement 1. Fgf18 receptor *Fgfr4* is predominantly expressed by myogenic cells in the LVP regions.

To identify perimysial-to-myogenic interactions that are functionally required for regulating myogenic fate, we decided to narrow our selection to the signaling molecules more specifically expressed by the perimysial fibroblasts. By checking the expression patterns of these signaling molecules in E13.5-E15.5 soft palate scRNAseq data, we found that *Lama4*, *Fgf18*, and *Dlk1* are the top three molecules more enriched in the perimysial fibroblasts than in other cells. This finding suggests they play a more specific role related to the myogenic site (Figure 5B). Among these three molecules, expression of *Lama4* and *Fgf18* was found to be reduced in the affected in *Osr2-Cre;Alk5^fl/fl^*late perimysial cells (Figure 5C and D). This suggests that TGF-β signaling could also regulate the expression of perimysial-derived *Lama4* and *Fgf18* during muscle development. *In vivo* expression of these two genes further showed that *Fgf18* expression is more restricted to the region of perimysial fibroblasts (Figure 5F) than that of *Lama4* (Figure 5G), similar to the TGF-β signaling activity; accordingly, we identified *Fgf18* as the most likely perimysial-to-myogenic signal regulated by TGF-β to support myogenesis.

To better understand the role of perimysial-myogenic signals during myogenesis, we further investigated the *in vivo* expression of *Fgf18* in perimysial cells during LVP development in control and *Osr2-Cre;Alk5^fl/fl^* mice. We found consistently enriched expression of *Fgf18* in the location of perimysial fibroblasts most adjacent to the myogenic cells between E14.5-E16.5 (Figure 5H-I and L-M). Consistent with the scRNAseq analysis, the expression of *Fgf18* in similar locations in the *Osr2-Cre;Alk5^fl/fl^* mice was reduced compared to their control counterparts (Figure 5J-K and N-O). Notably, the reduction of the myogenic cells was correlated with the loss of *Fgf18* expression patterns. In the *Osr2-Cre;Alk5^fl/fl^* soft palate at E14.5, a reduced number of *Myod1*+ myogenic cells was detectable in the location of the remnant *Fgf18* expression (Figure 5K) compared with the presence of myogenic cells in control (Figure 5I). Further, at E16.5, the *Myod1* expression became undetectable as *Fgf18* expression disappeared (Figure 5O) compared with the abundant presence of both signals in the control (Figure 5M). This strong correlation between the perimysial-fibroblast-derived *Fgf18* and *Myod1+* myogenic cells further supported the functional role of *Fgf18* in myogenesis. We therefore next evaluated the expression of Fgf receptors in the MyoD*+* myogenic cells to test whether myogenic cells could receive an Fgf18 signal. Consistent with CellChat predictions, late-perimysial Fgf18 signals were more likely to communicate with Fgfr1 and/or Fgfr4 in the myogenic cells, which did not express *Fgfr2* or *Fgfr3* (Figure 5—figure supplement 1B, C, F, and G). Though *Fgfr1* is expressed extensively in the palatal mesenchymal cells and weakly in myogenic cells (Figure 5—figure supplement 1A and E), *Fgfr4* showed more specific and stronger expression in the myogenic cells (Figure 5—figure supplement 1D and H). Thus, perimysial-fibroblast-derived Fgf18 may signal to Fgfr4 in the myogenic cells to orchestrate myogenic-specific functions.

### Smad2/3 and Creb5 cooperatively regulate Fgf18 signaling for pharyngeal muscle development

To investigate how TGF-β signaling serves as a key regulator of perimysial-to-myogenic communication, particularly Fgf18 signaling, we first explored whether Smad2/3 can directly bind to the promoter region of Fgf18 using JASPAR TF binding sites (TFBSs) predictions incorporated in the UCSC Genome Browser (Castro-Mondragon *et al*., 2022). Several genome tracks of TFBSs for Smad2/3 are present in the promoter region 2 kb upstream of the transcription start site (TSS) (Figure 6A). We focused on the binding motif (chr11:33098353-33098362) with the highest predicted binding score since it is most likely to be the binding site (Figure 6B). Using a Cut and Run assay for Smad2/3, we identified significantly greater enriched DNA fragments around the binding motif than IgG-enriched (Figure 6C), suggesting direct binding of Smad2/3 to this predicted TFBS in the *Fgf18* promoter region. Thus, *Fgf18* is a direct target for TGF-β signaling in the late perimysial cells.

**Figure 6.**
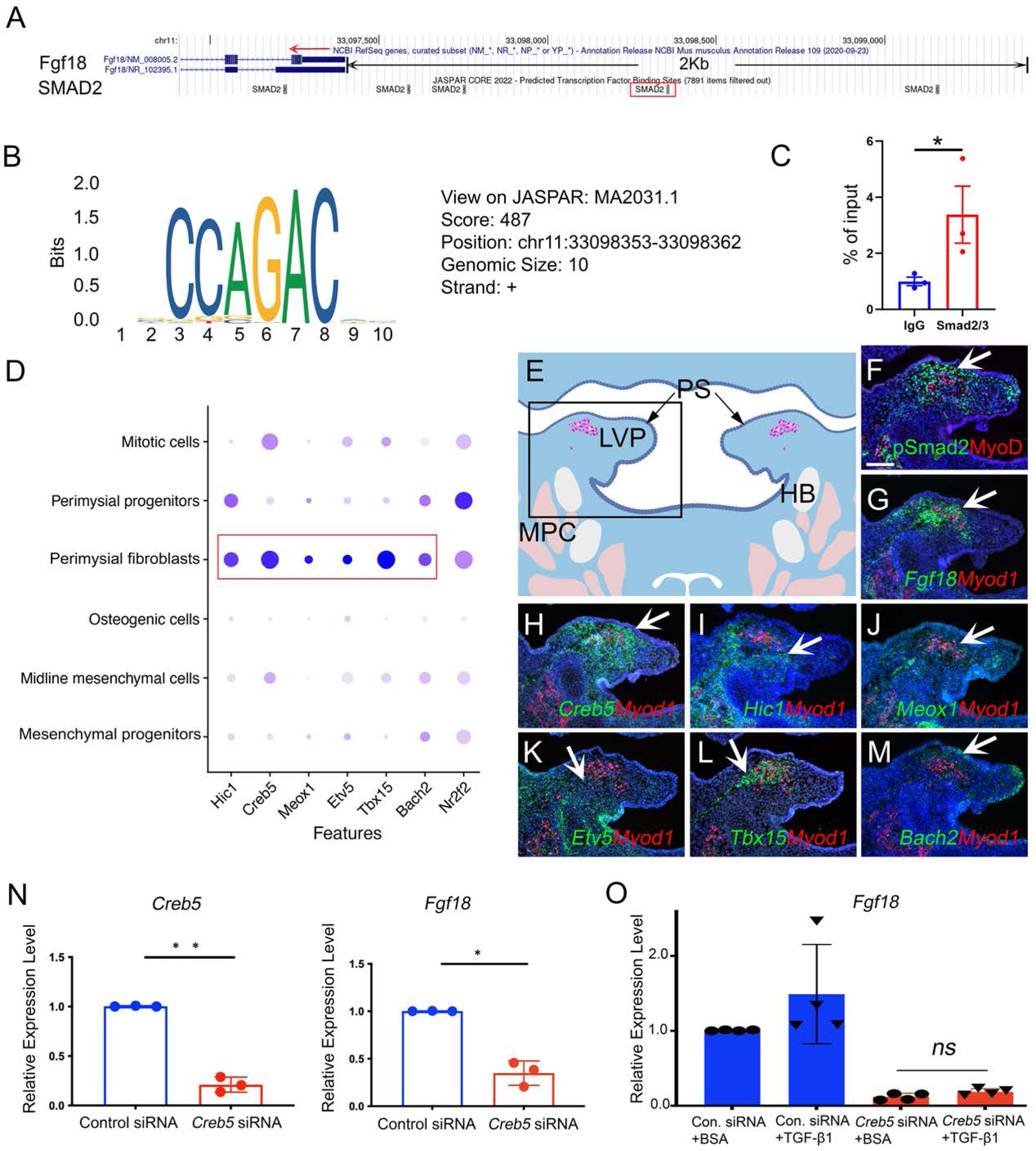
Perimysial fibroblast-specific regulon *Creb5*, identified by SCENIC analysis, cooperates with TGF-β signaling to regulate *Fgf18.* (A) UCSC binding prediction of SMAD2 binding motif to the promoter of *Fgf18* gene. The boxed area indicates the binding site with the highest score. (B) The predicted binding motif of the SMAD2 in the promoter region of *Fgf18* gene with the highest score. (C) Cut and Run analysis shows significantly more enriched SMAD2/3 binding to the promoter region of the *Fgf18* gene close to the predicted binding site than that of IgG in the soft palatal tissue of the control mice at E14.5. *, p-value ≤0.05. (D) Dotplot of perimysial fibroblast regulon expression pattern in individual cellular clusters of integrated scRNAseq from E13.5-E15.5 soft palatal tissue. Highlighted regulons are more enriched in the perimysial fibroblasts. (E) Schematic drawing of coronal section of the LVP region at E14.5. Boxed area in E indicates the region shown in F-M. The magenta and salmon colors represent the LVP myogenic cells and the other pharyngeal muscles, respectively. HB, hyoid bone; LVP, levator veli palatini; MPC, middle pharyngeal constrictor; PS, palatal shelves. (F) Immunofluorescence of MyoD (red) and pSMAD2 (green) in the coronal section of the LVP region at E14.5. (G-M) RNAScope *in situ* hybridization for *Myod1* (red) and *Fgf18* (green) (G), *Creb5* (green) (H), *Hic1* (green) (I), *Meox1* (green) (J), *Etv5* (green) (K), *Tbx15* (green) (L) or *Bach2* (green) (M) in coronal sections of the LVP region at E14.5. Arrows indicate positive signals. (N) qPCR analysis of *Creb5* and *Fgf18* expression after *Creb5* siRNA treatment on E14.5 soft palatal cell culture compared with the control siRNA. **, p≤0.01; * p≤0.05. (O) qPCR analysis of *Fgf18* expression following *Creb5* siRNA treatment combined with 5 ng/ml TGF-β1 or BSA on E14.5 soft palatal cell culture, compared with the control siRNA. N=4. ns, not significant. Scale bar in F indicates 100 μm in F-M. Figure 6-Source data 1 for 6C Figure 6-Source data 2 for 6N Figure 6-Source data 3 for 6O This article includes the following figure supplements for figure 6: Figure supplement 1. SCENIC analysis identified individual cell type-specific regulons from E13.5-E15.5 soft palatal scRNAseq data.Figure supplement 2. TGF-β signaling, *Creb5*, and *Fgf18* are expressed in a similar region of the perimysial cells of the masseter at E13.5.

The biological effects of TGF-β signaling are contextual (Morikawa *et al*., 2016). One mechanism for cell-type-specific response to TGF-β signaling is through the cooperation with cell-type-specific master transcription factors, which enable TGF-β signaling to specifically affect the cell-type-specific genes bound by these master regulators (Mullen *et al*., 2011). To identify master regulators that help TGF-β signaling activate a perimysial-specific response in palatal mesenchymal cells, we performed Single-Cell Regulatory Network Interference and Clustering (SCENIC) analysis using E13.5-E15.5 soft palate scRNAseq data (Aibar *et al*., 2017; Han *et al*., 2021). We identified that gene regulatory networks (GRNs) mediated by *Hic1*, *Creb5, Meox1*, *Etv5*, *Tbx15*, *Bach2*, and *Nr2f2* are most associated with late-perimysial cells (Figure 6— figure supplement 1). The expression patterns of these genes in scRNAseq data indicated that *Hic1*, *Creb5, Meox1, Etv5, Tbx15, and Bach2* are more enriched and specific to the perimysial fibroblasts than other palatal mesenchymal populations, and are thus more likely to be associated with the late-perimysial specific cell behaviors (Figure 6D). *In vivo* expression patterns further validated that, among these regulatory genes (Figure 6H-M), *Creb5* has the most specific expression pattern in the late perimysial cells (Figure 6H), similar to that of *Fgf18* and TGF-β signaling activity (Figure 6F and G). This overlapping expression further indicated that *Creb5* and TGF-β signaling may function cooperatively to activate perimysial-specific signaling genes such as *Fgf18*. Consistent with this observation, reduction of *Creb5* in E14.5 soft palate mesenchymal cell cultures led to reduction of *Fgf18* (Figure 6N), indicating that Creb5 can regulate *Fgf18* expression. Furthermore, TGF-β1 treatment could not restore the *Fgf18* expression after *Creb5* reduction (Figure 6O), suggesting that TGF-β signaling requires Creb5 to activate downstream targets like *Fgf18* in the late perimysial cells. Furthermore, we also found that TGF-β signaling activity (pSmad2), *Creb5* and *Fgf18* are also expressed in a similar pattern in the perimysial cells during the development of the other pharyngeal muscles, such as the first pharyngeal arch-derived masseter muscle (Figure 6—figure supplement 2), confirming the possibility that this TGF-β signaling and Creb5 cooperation in regulating perimysial signals, which we identified using the soft palatal muscle model, can plausibly be extended to other craniofacial muscles.

### Fgf18 reduction in perimysial cells leads to reduced myogenic cells in the LVP region of *Osr2-Cre;Fgf18^fl/fl^* mice while Fgf18 restoration partially rescues myogenic defects in *Osr2- Cre;Alk5^fl/fl^* samples

To test whether perimysial-derived Fgf18 is functionally required for myogenesis, we generated *Osr2-Cre;Fgf18^fl/fl^*mice with conditional knockout of *Fgf18* in the palatal mesenchymal cells. Unlike in the *Osr2-Cre;Alk5^fl/fl^* mice, loss of *Fgf18* in the *Osr2-Cre;Fgf18^fl/fl^* mice did not result in any obvious palatal shelf formation defects (Figure 7B and D) compared to the control mice (Figure 7A and C) at birth, consistent with recent studies demonstrating that Fgf18 is not required for the development of mesenchymal cells themselves in the hard palate (Xu *et al*., 2016; Yue *et al*., 2021). In contrast, the sizes of the soft palatal muscles of the *Osr2-Cre;Fgf18^fl/fl^* mice appeared to be smaller (Figure 7F and H) than those of controls (Figure 7E and G), which was especially obvious in the posterior part of LVP (Figure 7G and H). This was probably due to the difference in muscle fiber organization in the anterior and posterior parts of the LVP. The LVP consists of muscle fibers oriented both parallel and perpendicularly to the section plane in the anterior region (Figure 7I) and predominantly perpendicularly in the posterior region (Figure 7K). While all the fibers appear more sparse in the anterior and posterior LVP of the *Osr2- Cre;Fgf18^fl/fl^* mice (Figure 7J and L), the perpendicular fiber numbers are significantly reduced (Figure 7J). To further analyze the progression of muscle defects, we assessed the soft palatal and myogenic phenotypes from E14.5 onwards (Figure 7—figure supplement 1). While the MyoD+ myogenic cells appeared comparable in the control and *Osr2-Cre;Fgf18^fl/fl^* LVP at E14.5 (Figure 7—figure supplement 1A-D), the myogenic cells in the *Osr2-Cre;Fgf18^fl/fl^*mice started to appear progressively reduced compared to the control, and this difference became more apparent at E16.5 (Figure 7—figure supplement 1E-H), confirming the role of Fgf18 in late-stage myogenesis. We also confirmed that the loss of functional Exon1C of *Fgf18* is efficient in the *Osr2-Cre;Fgf18^fl/fl^* soft palatal shelves (Figure 7—figure supplement 1I-L). Taken together, these findings suggest that Fgf18 may not be required for determining the fate of palatal mesenchymal cells themselves, but rather may serve as a paracrine molecule for the myogenic cell development.

**Figure 7.**
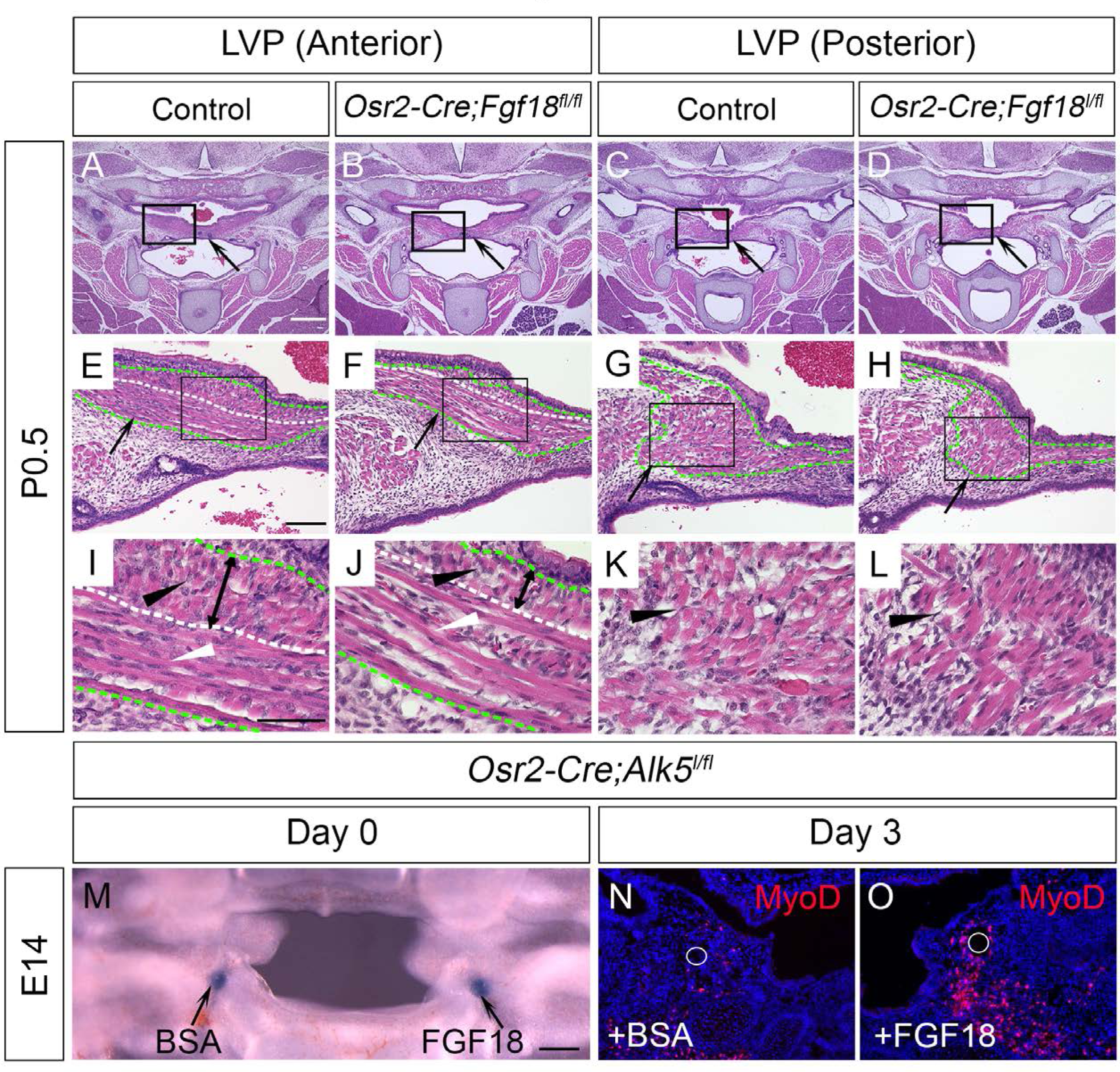
Myogenic cells are reduced in the LVP region of *Osr2-Cre;Fgf18^fl/fl^* mice, while Fgf18 increases MyoD+ myogenic cells in *Osr2-Cre;Alk5^fl/fl^* soft palate slice cultures. (A-L) H&E staining in coronal sections of LVP region at P0.5 from control (A, C, E, G, I, and K) and *Osr2-Cre;Fgf18^fl/fl^* mice (B, D, F, H, J, and L). Boxed areas in A-D and E-H are enlarged in E-H, and I-L, respectively. Black arrows in A-D point to the palatal shelf and the LVP in E-H. Green dotted line outlined the LVP in E-H. Black and white triangles point to perpendicular muscle fibers in I-L and parallel fibers in I-J. The white dotted line indicates the boundaries of perpendicular and parallel fibers in E, F, I and J. Double-ended arrows indicate the thickness of perpendicular fibers in I and J. (M) A 300 μm coronal slice of the LVP region at E14 from *Osr2- Cre;Fgf18^fl/fl^* mouse for slice culture following bead implantation. Arrows point to BSA- or FGF18-treated bead. (N-O) Immunofluorescence of MyoD (red) in the coronal section of LVP region from *Osr2-Cre;Fgf18^fl/fl^* mouse cultured for 3 days with BSA bead (N) and FGF18 bead (O). White circles indicate the location of the BSA beads in N and FGF18 beads in O. Scale bars in A, E, I and M indicate 500 μm in in A-D, 100 μm in E-H, 50 μm in I-L, and 100 μm in M-O, respectively. This article includes the following figure supplement for figure 7: Figure supplement 1. *Osr2-Cre;Fgf18^fl/fl^* mice exhibit myogenic defects during LVP development.

To test whether Fgf18 treatment could restore some of the severe muscle formation defects after following the loss of TGF-β signaling in *Osr2-Cre;Alk5^fl/fl^* mice, we optimized an organ culture system (Alfaqeeh and Tucker, 2013; Humpel, 2015) to culture 300 μm-thick slices of embryonic head tissues containing the LVP region. This enabled the maintenance of the three-dimensional structure of the pharyngeal region and allowed for efficient nutrient penetration and bead implantation (Figure 7M). Using this culture system, we found that FGF18 bead treatment in the soft palatal shelves led to increased MyoD+ cells compared with BSA beads in *Osr2-Cre;Alk5^fl/fl^* slice cultures (Figure 7N and O). This suggests that Fgf18 is one of the key regulators of perimysial-to-myogenic signaling affected in the LVP in the *Osr2-Cre;Alk5^fl/fl^*mice. Taken together, our studies identified that TGF-β signaling interacts with perimysial regulator Creb5 to specify the perimysial-to-myogenic signaling, such as Fgf18, in individual pharyngeal muscle development.

## Discussion

In the later stage of pharyngeal myogenesis, myogenic progenitors migrate into discrete myogenic sites to form individual muscle anlagen anatomically resembling their adult counterparts (Noden and Francis-West, 2006; Sambasivan *et al*., 2011; Shih *et al*., 2008; Ziermann *et al*., 2018). How these fine-tuned individual muscles form the right morphology at the right location in order to perform their physiological functions remains a question of interest. Here, using the pharyngeal muscle LVP as a model, our study showed that an important aspect of this mechanism is the establishment of a distinct pro-myogenic perimysial subdomain adjacent to the myogenic site. The cells in this domain have unique identities and regulatory mechanisms distinct from the rest of connective tissue cell populations within the palatal mesenchyme, enabling perimysial cells to uniquely provide various pro-myogenic signals and define specific myogenic sites to support region-specific myogenesis. Using unbiased screening with scRNAseq analysis combined with mouse genetic approaches, we identified TGF-β signaling as a predominant and specific regulator for perimysial cell fate determination during this stage, and perimysial-specific transcriptional factor Creb5 assists TGF-β signaling to achieve functional specificity in supporting pharyngeal myogenesis via pro-myogenic signals such as Fgf18.

In early myogenesis, CNC cells have also been shown to induce myogenic differentiation by secreting both BMP and Wnt inhibitors to antagonize the dorsal neural tube-derived BMP and Wnt signaling molecules that repress craniofacial skeletal muscle formation (Tzahor *et al*., 2003). In the later developmental stages, when the CNC-derived palatal mesenchymal cells are more differentiated and patterned into distinguishable functional domains, we have the optimal opportunity to explore a specific regulatory mechanism of myogenesis. We found that during the LVP development, the late perimysial population (perimysial fibroblasts) with the most active TGF-β signaling is also distributed according to a muscle-specific pattern in both the fourth pharyngeal arch-derived LVP and first pharyngeal arch-derived masseter muscle; and this TGF-β signaling function is required for the formation of the muscle analgen of both the LVP and masseter (Han *et al*., 2014). Similarly, in limb muscle development, a group of pre-patterned *Tcf4*+ limb mesodermal cells also pre-determines the basic pattern of the muscles (Kardon *et al*., 2003). This indicates a potentially universal mechanism in which a pre-defined perimysial domain distinct from the rest of the connective tissues is required to establish specific myogenic sites that allow for proper muscle analge specification in late developmental stages.

In late developmental stages, myogenic progenitors are also recruited to specific locations to form new muscles. By investigating perimysial-to-myogenic communication in this study, we identified pro-myogenic signaling from the neighboring late perimysial cells (perimysial fibroblasts). It thus appears that these embryonic myogenic progenitors may also require “embryonic niches” to support their contribution to muscle development at specific myogenic sites. In adults, muscle stem cells - satellite cells - reside in specific niches, and their ability to potentiate muscle repair and regeneration is also supported by signals from these niches (Andersen *et al*., 2013; Relaix *et al*., 2021). By comparing the “adult muscle stem cell niche” with the “embryonic niche,” more molecular and cellular similarities can be further identified. For example, several perimysial-derived signaling pathways for the embryonic myogenic progenitors are also required for satellite cell fate regulation after birth (Bigas and Espinosa, 2016; Farin *et al*., 2016; Stantzou *et al*., 2017). In addition, while *Hic1*+ mesenchymal progenitors in the adult niche coordinate various aspects of skeletal muscle regeneration (Scott *et al*., 2019), they are also inferred to be regulators of the perimysial cells of the embryonic niche during development. These similarities suggest potentially conserved mechanisms between embryonic muscle development and adult muscle repair/regeneration, consistent with the clinical observation that patients with disrupted soft palatal muscle development also exhibit impaired differentiation and a reduced number of satellite cells (Monroy *et al*., 2012). Thus, the mechanism we identified during development could also be potentially useful for promoting muscle growth after birth.

Craniofacial anomalies such as DiGeorge syndrome, as well as other syndromic and non-syndromic forms of cleft lip and palate, often affect pharyngeal muscle formation and lead to difficulties in eating, facial expression, speaking, and swallowing (Kelly *et al*., 2004; Kernahan *et al*., 1984; Monroy *et al*., 2012; Scambler, 2000). While the identification of a detailed Pitx2- Tbx1-Msc-Tcf21 transcription factor regulatory network for the pharyngeal myogenic cells may influence the intrinsic regulatory network within the myogenic cells to promote pharyngeal muscle development (Buckingham and Rigby, 2014; Sambasivan *et al*., 2011), manipulation through paracrine signaling is a more likely mechanism. In this study, we identified a variety of potential pro-myogenic signaling molecules from perimysial-to-myogenic cell interaction analysis. A “chemokine/cytokine cocktail” may be needed to restore the muscle detects in our cleft soft palate animal model (*Osr2-Cre;Alk5^fl/fl^*). As a proof of concept, we found partial improvement of the muscle defects after Fgf18 restoration and thus confirmed the potential of restoring pharyngeal muscle defects with the combination of pro-myogenic factors we identified in the study. Moreover, some of these molecules, including Fgf18 and Dlk1, have also been suggested to be involved in the development and regeneration of various muscles throughout the body including in the limb, diaphragm and tongue (Andersen *et al*., 2013; Ito *et al*., 2018; Mok *et al*., 2014; Yue *et al*., 2021), extending their potential application to defects in other muscles in the body.

In summary, our study has been able to decipher the molecular and cellular composition of the embryonic myogenic niche for the development of pharyngeal muscles and potentially other muscles in the body. Our work will contribute to a better understanding of the fine-tuned regulatory network of late-stage muscle morphogenesis and lead to the development of novel treatment strategies for regenerating/repairing muscle defects, particularly in infants with birth defects.

## Materials & Methods

### Key resources table

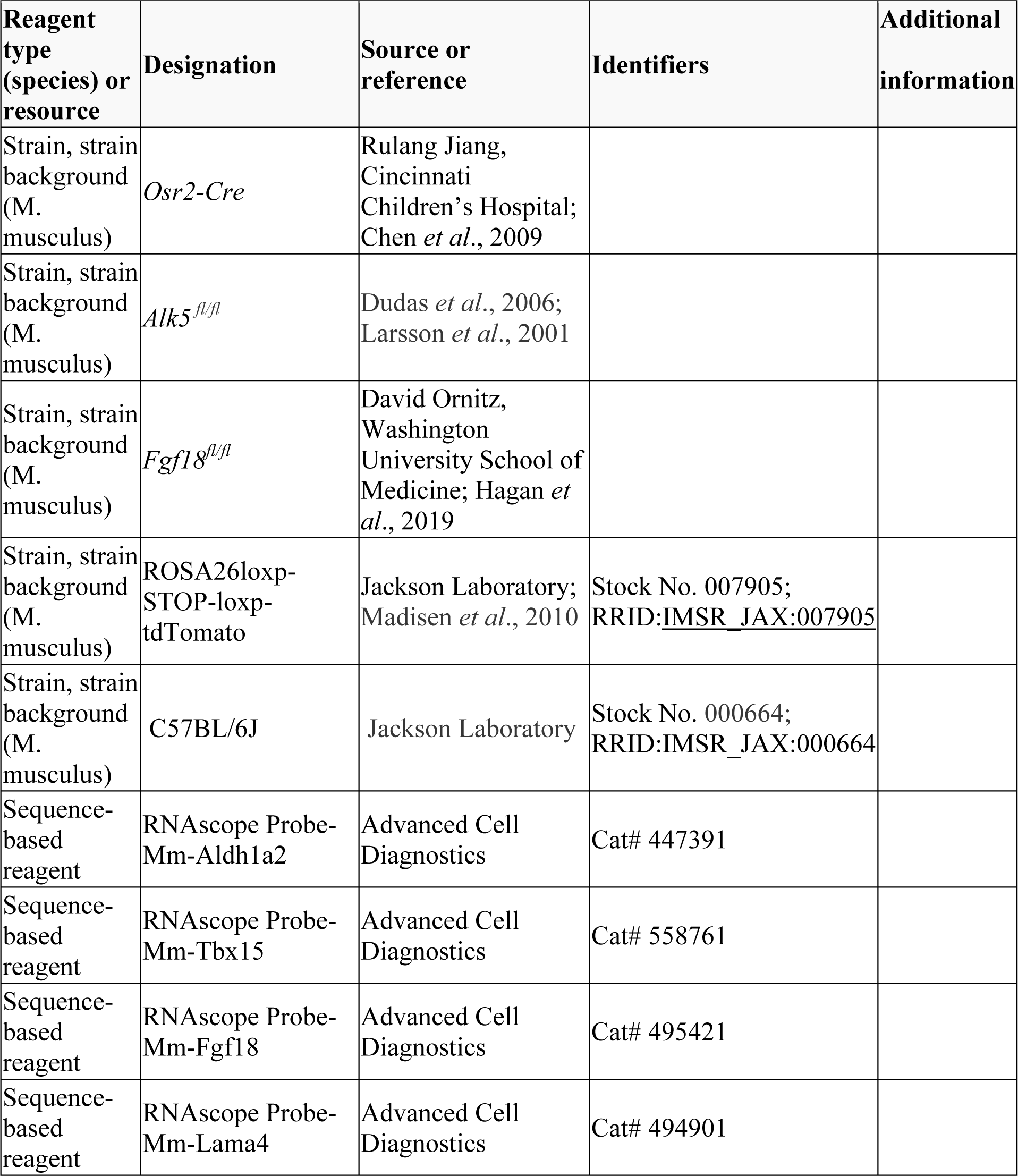
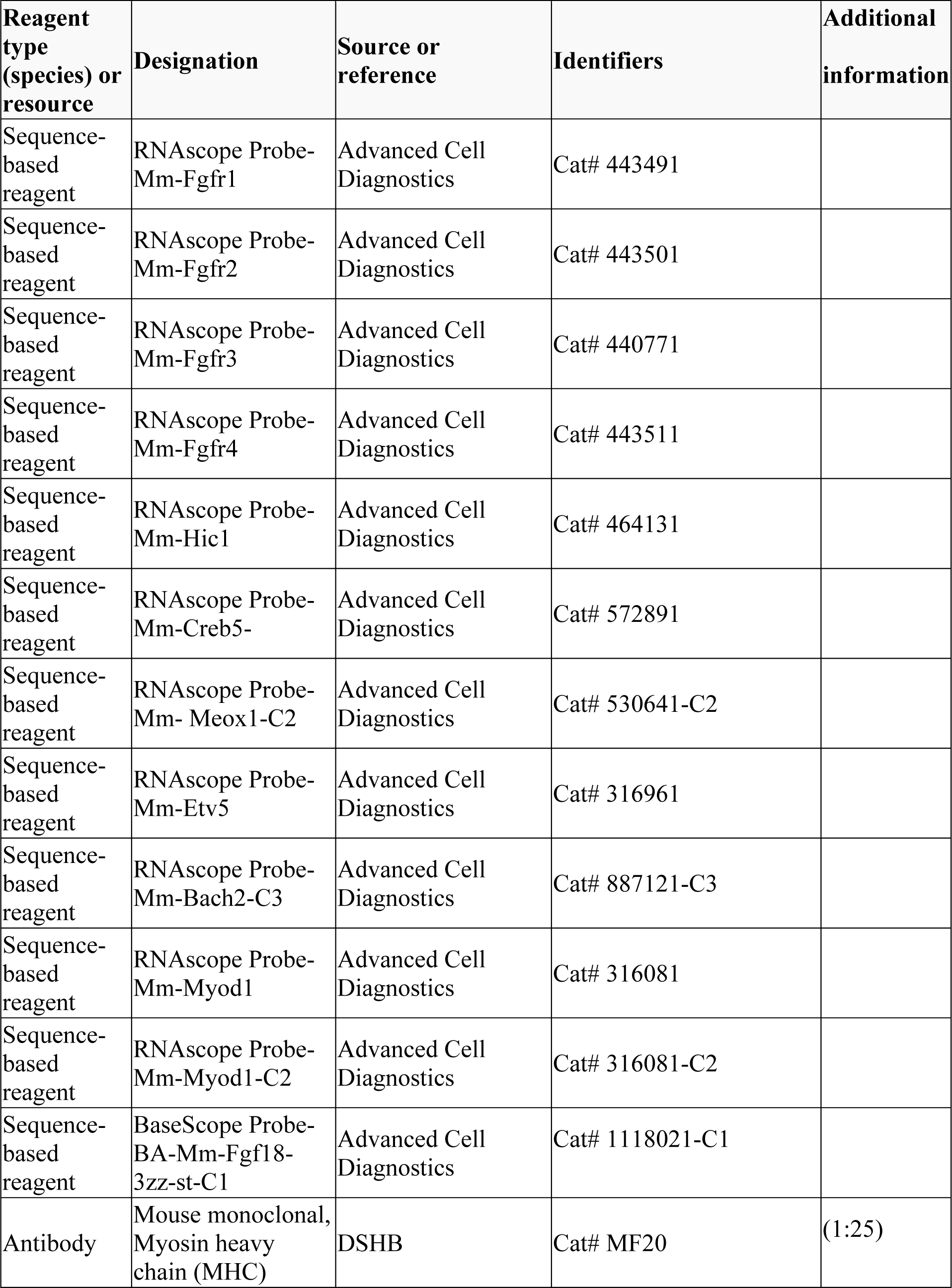
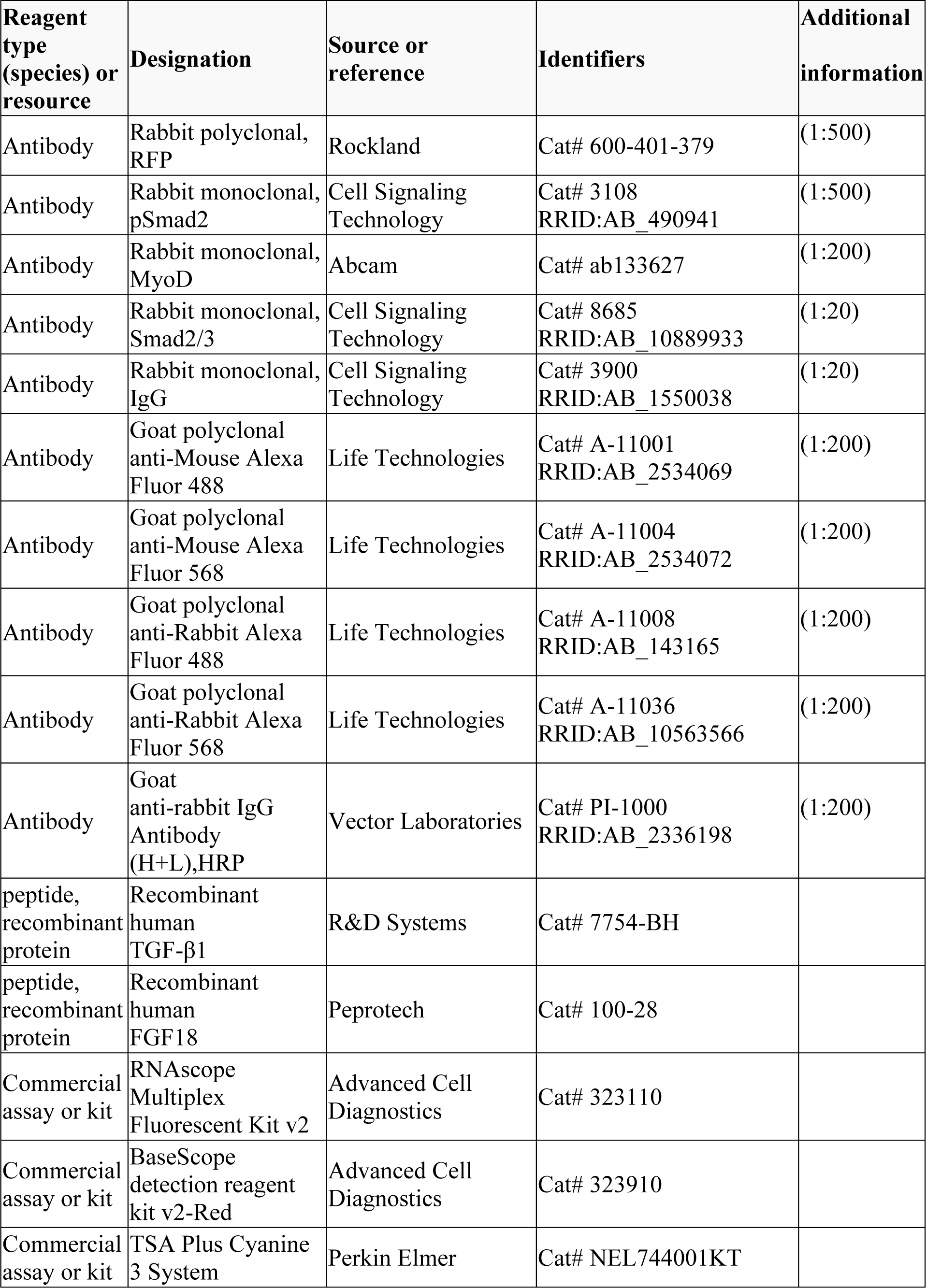
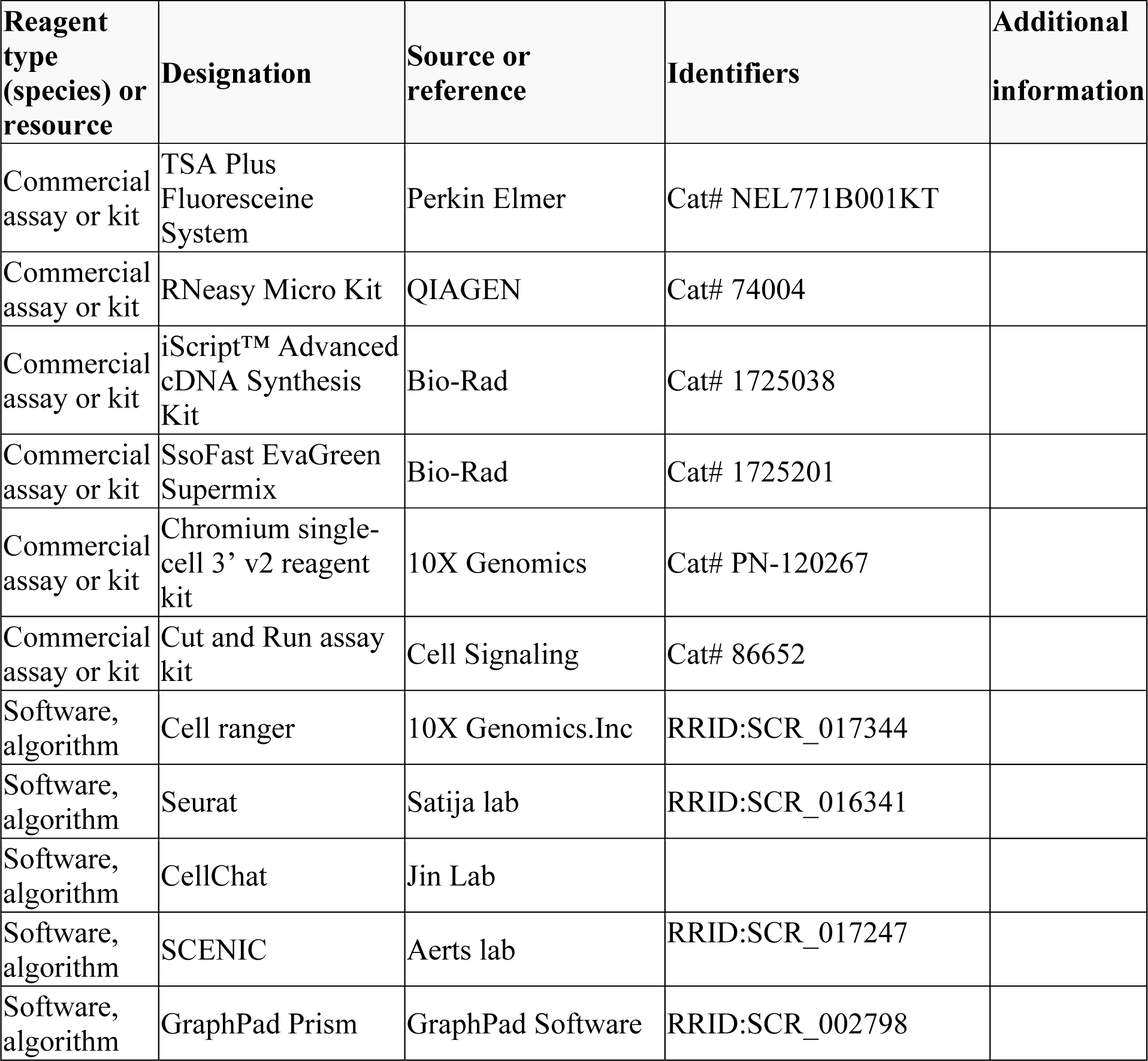

| Reagent type (species) or resource | Designation | Source or reference | Identifiers | Additional information |
| --- | --- | --- | --- | --- |
| Strain, strain background (M. musculus) | *Osr2-Cre* | Rulang Jiang, Cincinnati Children's Hospital; Chen *et al.*, 2009 |  |  |
| Strain, strain background (M. musculus) | *Alk5<sup>fl/fl</sup>* | Dudas *et al.*, 2006; Larsson *et al.*, 2001 |  |  |
| Strain, strain background (M. musculus) | *Fgf18<sup>fl/fl</sup>* | David Ornitz, Washington University School of Medicine; Hagan *et al.*, 2019 |  |  |
| Strain, strain background (M. musculus) | ROSA26loxP-STOP-loxP-tdTomato | Jackson Laboratory; Madisen *et al.*, 2010 | Stock No. 007905; RRID:IMSR_JAX:007905 |  |
| Strain, strain background (M. musculus) | C57BL/6J | Jackson Laboratory | Stock No. 000664; RRID:IMSR_JAX:000664 |  |
| Sequence-based reagent | RNAscope Probe-Mm-Aldh1a2 | Advanced Cell Diagnostics | Cat# 447391 |  |
| Sequence-based reagent | RNAscope Probe-Mm-Tbx15 | Advanced Cell Diagnostics | Cat# 558761 |  |
| Sequence-based reagent | RNAscope Probe-Mm-Fgf18 | Advanced Cell Diagnostics | Cat# 495421 |  |
| Sequence-based reagent | RNAscope Probe-Mm-Lama4 | Advanced Cell Diagnostics | Cat# 494901 |  |

| **Reagent type (species) or resource** | **Designation** | **Source or reference** | **Identifiers** | **Additional information** |
| --- | --- | --- | --- | --- |
| Sequence-based reagent | RNAscope Probe-Mm-Fgfr1 | Advanced Cell Diagnostics | Cat# 443491 |  |
| Sequence-based reagent | RNAscope Probe-Mm-Fgfr2 | Advanced Cell Diagnostics | Cat# 443501 |  |
| Sequence-based reagent | RNAscope Probe-Mm-Fgfr3 | Advanced Cell Diagnostics | Cat# 440771 |  |
| Sequence-based reagent | RNAscope Probe-Mm-Fgfr4 | Advanced Cell Diagnostics | Cat# 443511 |  |
| Sequence-based reagent | RNAscope Probe-Mm-Hic1 | Advanced Cell Diagnostics | Cat# 464131 |  |
| Sequence-based reagent | RNAscope Probe-Mm-Creb5- | Advanced Cell Diagnostics | Cat# 572891 |  |
| Sequence-based reagent | RNAscope Probe-Mm- Meox1-C2 | Advanced Cell Diagnostics | Cat# 530641-C2 |  |
| Sequence-based reagent | RNAscope Probe-Mm-Etv5 | Advanced Cell Diagnostics | Cat# 316961 |  |
| Sequence-based reagent | RNAscope Probe-Mm-Bach2-C3 | Advanced Cell Diagnostics | Cat# 887121-C3 |  |
| Sequence-based reagent | RNAscope Probe-Mm-Myod1 | Advanced Cell Diagnostics | Cat# 316081 |  |
| Sequence-based reagent | RNAscope Probe-Mm-Myod1-C2 | Advanced Cell Diagnostics | Cat# 316081-C2 |  |
| Sequence-based reagent | BaseScope Probe-BA-Mm-Fgf18-3zz-st-C1 | Advanced Cell Diagnostics | Cat# 1118021-C1 |  |
| Antibody | Mouse monoclonal, Myosin heavy chain (MHC) | DSHB | Cat# MF20 | (1:25) |
| Antibody | Rabbit polyclonal, RFP | Rockland | Cat# 600-401-379 | (1:500) |
| Antibody | Rabbit monoclonal, pSmad2 | Cell Signaling Technology | Cat# 3108<br>RRID:AB_490941 | (1:500) |
| Antibody | Rabbit monoclonal, MyoD | Abcam | Cat# ab133627 | (1:200) |
| Antibody | Rabbit monoclonal, Smad2/3 | Cell Signaling Technology | Cat# 8685<br>RRID:AB_10889933 | (1:20) |
| Antibody | Rabbit monoclonal, IgG | Cell Signaling Technology | Cat# 3900<br>RRID:AB_1550038 | (1:20) |
| Antibody | Goat polyclonal anti-Mouse Alexa Fluor 488 | Life Technologies | Cat# A-11001<br>RRID:AB_2534069 | (1:200) |
| Antibody | Goat polyclonal anti-Mouse Alexa Fluor 568 | Life Technologies | Cat# A-11004<br>RRID:AB_2534072 | (1:200) |
| Antibody | Goat polyclonal anti-Rabbit Alexa Fluor 488 | Life Technologies | Cat# A-11008<br>RRID:AB_143165 | (1:200) |
| Antibody | Goat polyclonal anti-Rabbit Alexa Fluor 568 | Life Technologies | Cat# A-11036<br>RRID:AB_10563566 | (1:200) |
| Antibody | Goat anti-rabbit IgG Antibody (H+L),HRP | Vector Laboratories | Cat# PI-1000<br>RRID:AB_2336198 | (1:200) |
| peptide, recombinant protein | Recombinant human TGF-β1 | R&D Systems | Cat# 7754-BH | |
| peptide, recombinant protein | Recombinant human FGF18 | Peptotech | Cat# 100-28 |  |
| Commercial assay or kit | RNAscope Multiplex Fluorescent Kit v2 | Advanced Cell Diagnostics | Cat# 323110 |  |
| Commercial assay or kit | BaseScope detection reagent kit v2-Red | Advanced Cell Diagnostics | Cat# 323910 |  |
| Commercial assay or kit | TSA Plus Cyanine 3 System | Perkin Elmer | Cat# NEL744001KT |  |
| Commercial assay or kit | TSA Plus Fluoresceine System | Perkin Elmer | Cat# NEL771B001KT |  |
| Commercial assay or kit | RNeasy Micro Kit | QIAGEN | Cat# 74004 |  |
| Commercial assay or kit | iScript™ Advanced cDNA Synthesis Kit | Bio-Rad | Cat# 1725038 |  |
| Commercial assay or kit | SsoFast EvaGreen Supermix | Bio-Rad | Cat# 1725201 |  |
| Commercial assay or kit | Chromium single-cell 3' v2 reagent kit | 10X Genomics | Cat# PN-120267 |  |
| Commercial assay or kit | Cut and Run assay kit | Cell Signaling | Cat# 86652 |  |
| Software, algorithm | Cell ranger | 10X Genomics.Inc | RRID:SCR_017344 |  |
| Software, algorithm | Seurat | Satija lab | RRID:SCR_016341 |  |
| Software, algorithm | CellChat | Jin Lab |  |  |
| Software, algorithm | SCENIC | Aerts lab | RRID:SCR_017247 |  |
| Software, algorithm | GraphPad Prism | GraphPad Software | RRID:SCR_002798 |  |

### Animal studies

The *Osr2-Cre* (gift from Rulang Jiang, Cincinnati Children’s Hospital; Chen *et al*., 2009), tdTomato conditional reporter (JAX#007905, Jackson Laboratory; Madisen *et al*., 2010), *Alk5^fl/fl^* (Dudas *et al*., 2006; Larsson *et al*., 2001), *Fgf18 ^fl/fl^* (gift from David Ornitz, Washington University School of Medicine; Hagan *et al*., 2019), and C57BL/6J (JAX#000664, Jackson Laboratory) mouse lines have all been described previously. To generate *Osr2- Cre;tdTomato^fl/fl^* mice, we crossed *Osr2-Cre* male mice with *tdTomato^fl/fl^* female mice. To generate *Osr2-Cre;Alk5^fl/fl^* mice, we crossed *Osr2-Cre;Alk5^fl/+^* male mice with *Alk5^fl/fl^* female mice. To generate *Osr2-Cre;Fgf18^fl/fl^*mice, we bred male *Osr2-Cre;Fgf18^fl/+^* mice with female *Fgf18^fl/fl^* mice. All mice were genotyped using genotyping primers as previously reported. The embryonic samples and newborn pups were collected and used for analysis without consideration of sex. All animal handling followed federal regulation and was performed with the approval of the Institutional Animal Care and Use Committee (IACUC) at the University of Southern California documented under protocol numbers 9320 and 11765.

### Tissue processing

Embryonic and newborn mouse heads were dissected and fixed in 10% formalin (HT501128, MilliporeSigma) overnight at room temperature following decalcification in 10% EDTA depending on the stage. For paraffin sections, samples were processed in serially ascending concentrations of ethanol solution at room temperature followed by xylene and paraffin wax at 60°C, then embedded in paraffin wax and sectioned at 8μm using a microtome (RM2255, Leica). Deparaffinized sections were stained with Hematoxylin and Eosin (H&E). For cryosections, samples were dehydrated in 15% sucrose/PBS solution followed by 30% sucrose/50% Tissue-Tek OCT compound (4583, Sakura). Samples were embedded in the OCT compound and frozen on a block of dry ice. Embedded samples were then cryosectioned at 8 μm using a cryostat (CM13050S, Leica).

### Immunostaining

Sections were antigen-retrieved for 10 minutes in preheated antigen unmasking solution (H-3300, Vector Laboratories) followed by 10 minutes of incubation with 1% Triton X in PBS (T8787, Sigma Aldrich). Sections were then incubated with 1% blocking reagent (PerkinElmer, FP1012) for 1 hour and then primary antibody overnight at 4°C. The sections were then incubated for 2 hours at room temperature with Alexa Fluor 488 or 568-conjugated secondary antibodies or HPR conjugated antibodies (Vector). For TSA-based immunofluorescent staining, the sections were further incubated with 1:200 TSA Plus FITC or Cy3 reagents for 3-5 minutes of signal development (NEL774001KT or NEL771B001KT, Akoya Bioscience). Sections were then counterstained with 4′,6-diamidino-2-phenylindole (DAPI; D9542, Sigma-Aldrich) and mounted with Fluoro-Gel (17985-10, EMS). The sections were washed in phosphate-buffered saline (PBS) with 0.1% Tween 20 (P1379, Sigma) between incubations. The primary and secondary antibodies used in this study were pSmad2 (3108, Cell signaling, 1:500 with TSA), MyoD (ab133627, Abcam; 1:200 with TSA), RFP (600-401-379, Rockland, 1:500 with TSA), myosin heavy chain (MHC) (MF20, DSHB, 1:25), Alexa Fluor 488 anti-mouse (A11001, Thermo fisher Scientific, 1:200), Alexa Fluor 568 anti-mouse (A11004, Thermo Fisher Scientific, 1:200), Alexa Fluor 488 anti-rabbit (A11008, Thermo Fisher Scientific, 1:200); Alexa Fluor 568 anti-rabbit (A11036, Thermo Fisher Scientific, 1:200), and anti-rabbit HRP (PI-1000, Vector Laboratories, 1:200).

### RNAscope *in situ* hybridization (ISH) assay

Tissue sections were air-dried at 60°C for 1 hour to overnight. RNAScope multiplex fluorescent reagent kit v2 (323100, Advanced Cell Diagnostics) and BaseScope detection reagent kit v2-Red (323910, Advanced Cell Diagnostics) were used for *in situ* hybridization according to the manufacturer’s instructions. RNAScope probes from Advanced Cell Diagnostics used in this study were *Aldh1a2* (447391), *Tbx15* (558761), *Fgf18* (495421), *Fgfr1* (443491), *Fgfr2* (443501s), *Fgfr3* (440771), *Fgfr4* (443511), *Lama4* (494901), *Hic1* (464131), *Creb5* (572891), *Meox1* (530641-C2), *Etv5* (316961), *Bach2* (887121-C3), *Myod1* (316081 or 316081-C2), and *Fgf18-Exon1C* (1118021-C1).

### Single-cell RNAseq

Soft palate tissue (dissected from the posterior third of the palatal region) from E14.5 control and *Osr2-Cre;Alk5^fl/fl^* embryos was digested and dissociated into single-cell suspension with TrypLE Express enzyme (12605010, Thermo Fisher Scientific) at 37°C with shaking at 600 rpm for 15 minutes using a Thermomixer (2231000269, Thermo Fisher Scientific). Single-cell suspension was loaded into the 10X Chromium system and prepared for single-cell library construction using the 10X Genomics Chromium single-cell 3’ v2 reagent kit (PN-120267, 10X Genomics) according to the manufacturer’s protocol. Sequencing was performed on Novaseq 6000 (Illumina). Library quality control, sequence alignment, and read counts were analyzed using Cell Ranger 4.0.0. Two control samples were combined using the function *cellranger aggr* so that the combined control sample had comparable cell numbers to the sample from *Osr2- Cre;Alk5^fl/fl^*. For each sample, raw read counts from every single cell were analyzed to identify cell clusters and variably expressed genes in each cluster using the Seurat R package (Hao *et al*., 2021) as previously described (Han *et al*., 2021). Seurat was also used to combine the E14.5 control and *Osr2-Cre;Alk5^fl/fl^* embryos to perform integration analysis as previously described (Han *et al*., 2021). RunPCA and RunUMAP visualization were used for downstream analysis and visualization. The integrated Seurat object combining E13.5-E15.5 control soft palates generated in our previous study was used to analyze gene regulatory network inference and ligand-receptor interactions. Gene regulatory network inference was performed using the R package SCENIC (Aibar *et al*., 2017). Transcription factors for each cell population in the palate were identified using GENIE3 and compiled into regulons, then subjected to cis-regulatory motif analysis. Regulon activity was then scored using AUCell. CellChat (Jin *et al*., 2021) was used to identify the potential ligand-receptor interactions between cell populations in the soft palate. Pre- processing functions (identifyOverExpressedGenes, identifyOverExpressedInteractions and projectData) and core functions (computeCommunProb, computeCommunProbPathway and aggregateNet with standard parameters) were applied along with other functions (netVisual_circle, netVisual_bubble, netAnalysis_signalingRole_heatmap netAnalysis_dot, and netAnalysis_river) to determine the senders and receivers.

### Cut and Run assay

Soft palatal tissue of E14.5 C57BL/6J mouse embryos was dissected and digested using Multi Tissue Dissociation Kit 3 (130-110-204, Miltenyi Biotec) at 37°C with shaking at 600 rpm for 15 minutes in a Thermomixer (2231000269, Thermo Fisher Scientific), followed by passing through 70 μm Pre-Separation Filters (130-110-204, Miltenyi Biotec) to achieve single-cell suspension solution. The single-cell suspension was processed using a Cut and Run assay kit (86652, Cell Signaling) following the manufacturer’s instructions. Five μl of Smad2/3 (8685, Cell Signaling) or IgG (3900, Cell Signaling) were used for each reaction. Standard protocol qPCR reaction was run using SsoFast EvaGreen Supermix (Bio-Rad, 172-5202) on a Bio-Rad CFX96 Real-Time Systems. The sequences of qPCR primers used to detect enriched DNA fragments near the Smad2 binding site at Chr11:33038353-33098362 were GGTGGGGTGACTCAACTGAA (forward) and TTGCGTGGCCTAAGGGTAAG (reverse).

### Cell culture and qPCR analysis

Soft palatal tissue of E14.5 C57BL/6J mouse embryos was dissected and digested using Multi Tissue Dissociation Kit 3 (130-110-204, Miltenyi Biotec) at 37°C with shaking at 600 rpm for 15 minutes in a Thermomixer (2231000269, Thermo Fisher Scientific), followed by passing through 70 μm Pre-Separation Filters (130-110-204, Miltenyi Biotec) to achieve single-cell suspension solution. The cells were seeded in 24-well plates (0.5 x 10^5^ cells/well) with collagen coating solution (125-50, MilliporeSigma) and cultured with 10% Fetal Bovine Serum (12662029, Thermo Fisher Scientific)/DMEM (10566016, Thermo Fisher Scientific)/1% Penicillin-Streptomycin (15140148, Thermo Fisher Scientific). For siRNA treatment, *Creb5* siRNA (siRNA ID:s107062, 4390816, Thermo Fisher Scientific) and control siRNA (4390844, Thermo Fisher Scientific) were delivered to the cells using Lipofectamine™ RNAiMAX Transfection Reagent (13778100, Thermo Fisher Scientific) according to manufacturer’s instructions. The cells were then treated with the *Creb5* siRNA and control siRNA for 48 hours before analysis. For TGFβ1 treatment, recombinant human TGFβ1 (7754-BH, R&D Systems) was used at 5 ng/ml for the TGFβ1 treatment group for 24 hours, while an equal volume of BSA (RB04, R&D Systems) was added to the control group for the same period. RNeasy Plus Micro Kit (Qiagen, 74034) was used to isolate RNA. After RNA isolation, cDNA was transcribed using the iScript™ Advanced cDNA Synthesis Kit (Bio-Rad, 1725038). qPCR reaction was run using SsoFast EvaGreen Supermix (Bio-Rad, 1725201) on a Bio-Rad CFX96 Real-Time System. Primer sequences used for qPCR reactions were obtained from PrimerBank (Wang *et al*., 2011) as follows: *Creb5* (Forward-AGGATCTTCTGCCGTCTTGAT; Reverse- GCGCAGCCTTCAGTCTCAT), *Fgf18* (Forward-CCTGCACTTGCCTGTGTTTAC; Reverse-TGCTTCCGACTCACATCATCT), and *Gapdh* (Forward-AGGTCGGTGTGAACGGATTTG; Reverse-TGTAGACCATGTAGTTGAGGTCA).

### Embryonic head slice cultures

Embryonic mouse heads were dissected and collected in complete BGJb medium: BGJb Medium (Thermo Fisher Scientific, 12591038) with 10% Fetal Bovine Serum (12662029, Thermo Fisher scientific), 0.1 mg/ml L-ascorbic acid (A4403, Sigma-Aldrich), and 1% Penicillin-Streptomycin (15140148, Thermo Fisher Scientific). Mouse heads were embedded on ice in a mixture of preheated 20% gelatin (G2500-500G, MilliporeSigma) in BGJB culture medium and sectioned to 300 μm slices using MicroSlicer Zero 1N (Ted Pella, Inc., 10111N). These slices were placed on cell culture inserts (PICM0RG50, Sigma Aldrich) which were inserted into 6-well culture plates and cultured at 37°C in a 5% CO_2_ incubator in complete BGJb medium. For bead implantation, Affi-Gel blue agarose beads (1537302, BioRad) were soaked in 1 μg/μl FGF18 (100-28, Peprotech) or BSA for one hour at 37°C (Xu *et al*., 2016) and then implanted into the E14 soft palatal shelves of *Osr2-Cre;Alk5^fl/fl^* slices. The samples were collected 3 days after bead implantation.

### Imaging

The immunofluorescent and RNAScope *in situ* hybridization fluorescent signals were captured using a Leica DMI3000 B research microscope. The brightfield images were captured using Keyence BZ-X710.

### Statistical analysis

Sample sizes were extrapolated from sample sizes of previously published studies. No randomization or blinding were performed. No samples were excluded from the analysis. N=3 (3 independent biological replications) for all analyses unless otherwise indicated. For qPCR analysis, more than 2 technical replicates were used for measuring each sample. GraphPad Prism was used for statistical analysis. All bar graphs display mean ± SEM. Independent two-tailed Student’s t-test was applied to assess statistical significance. The chosen level of significance for all statistical tests in this study was p ≤0.05.

### Data availability

The E14.5 control and *Osr2-Cre;Alk5^fl/fl^* soft palate single-cell RNAseq data generated in this study have been deposited in Gene Expression Omnibus (GEO) under accession code GSE203035. The single-cell RNAseq data for E13.5-E15.5 soft palate (Xia et al., 2021) used for analysis in this study have been deposited in GEO under accession code GSE155928.

## Acknowledgments

We thank Bridget Samuels for the critical reading of the manuscript and Giselle Mejia and Andrea Diaz for the schematic illustrations. We acknowledge USC Libraries Bioinformatics Service for assisting with data analysis. The bioinformatics software and computing resources used in the analysis were funded by the USC Office of Research and the Norris Medical Library. This study was supported by grants from the National Institute of Dental and Craniofacial Research, National Institutes of Health (R01 DE012711, U01 DE028729, and R90 DE022528).

---

**Figure 1—figure supplement 1.**
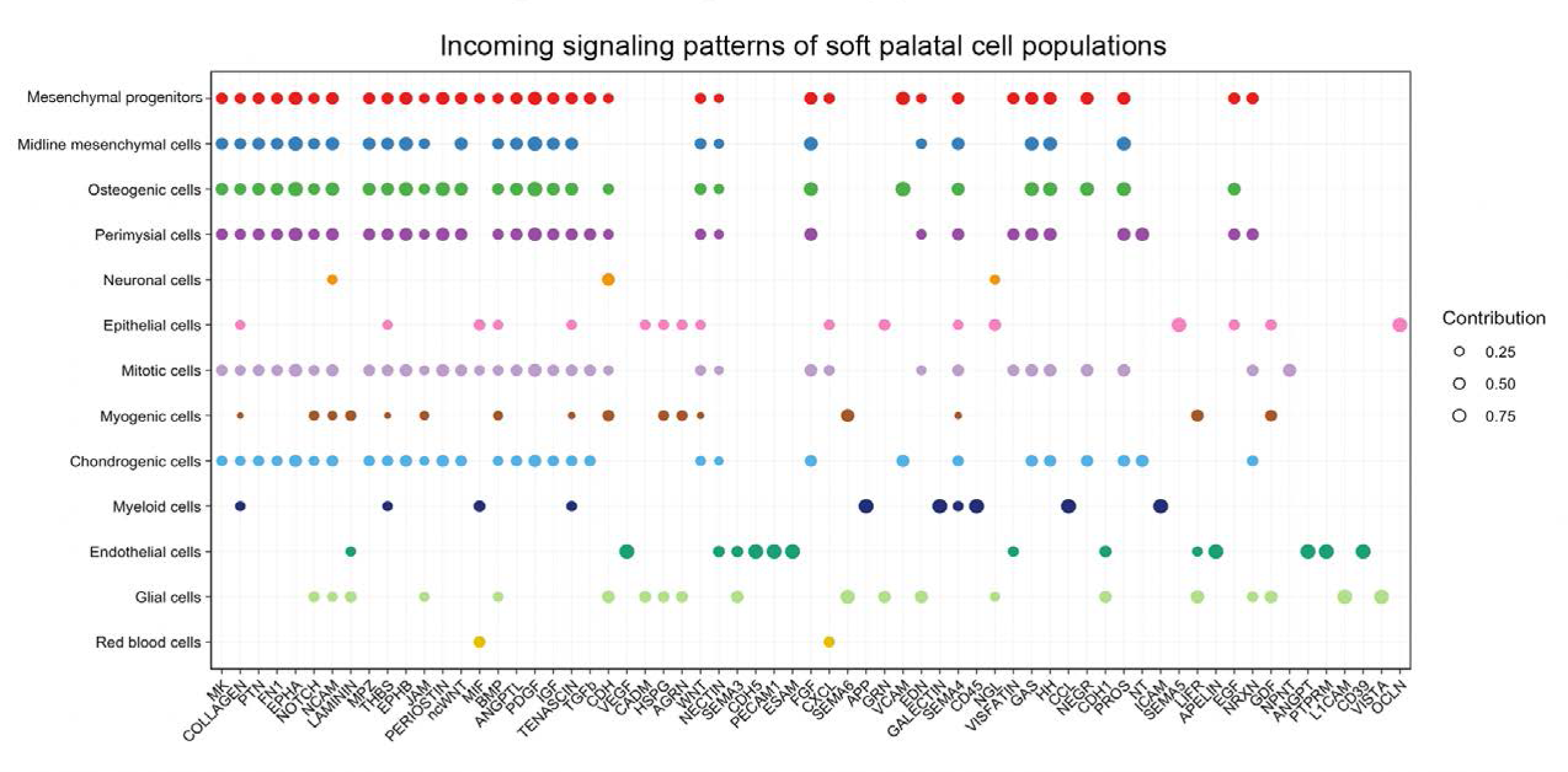
CellChat analysis of scRNAseq data identified incoming signaling pathways for all cell types in the soft palate. Bubble plot showing significant incoming signaling patterns for all the cell types of developing soft palate from E13.5-E15.5. Different colors in the plot represent different cell groups. Sizes in the bubble plot are proportional to the communication probability.

**Figure 3—figure supplement 1.**
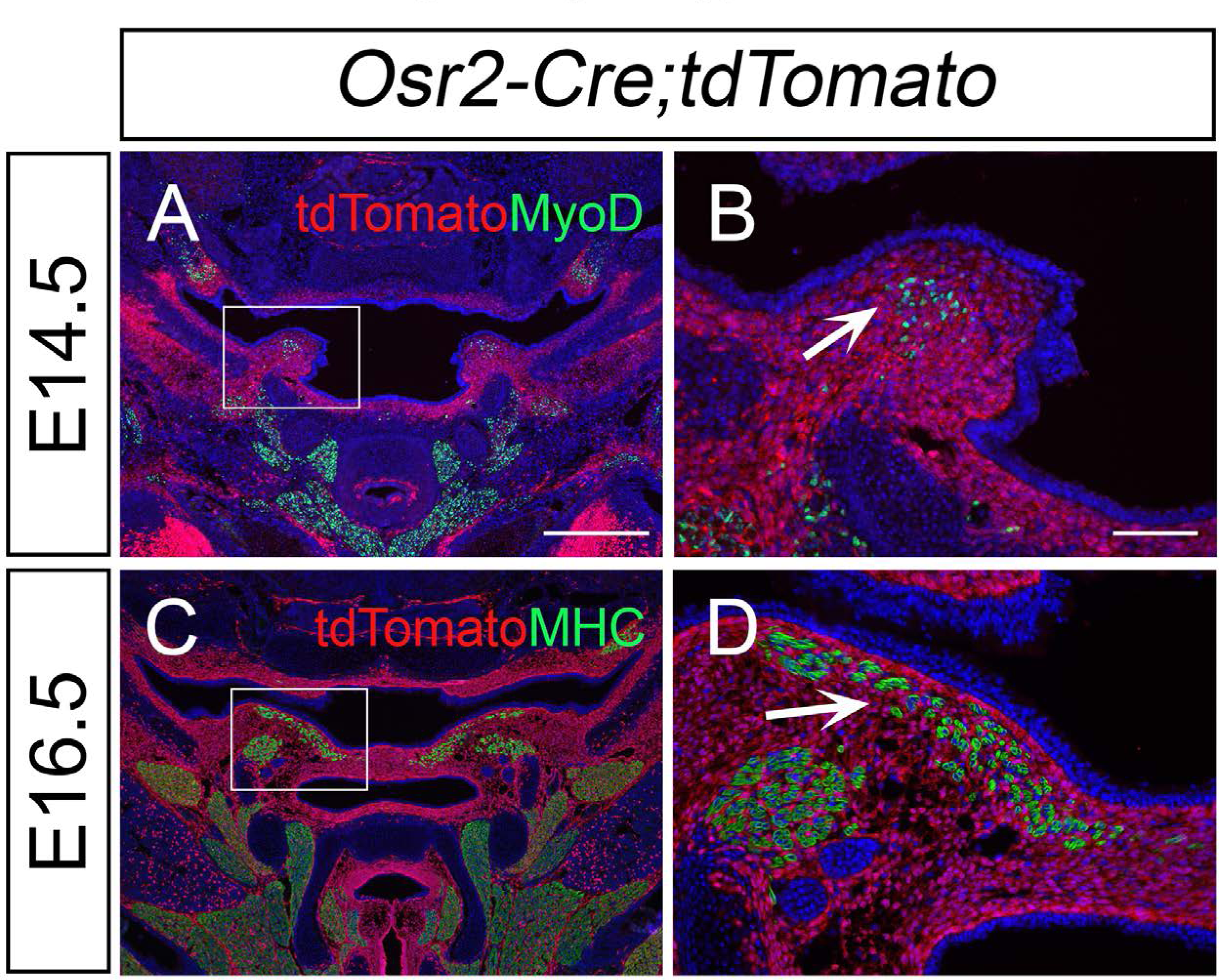
*Osr2-Cre* targets palatal mesenchymal cells surrounding soft palatal myogenic cells. (A-D) Immunofluorescence of myogenic marker MyoD (green) or MHC (green) and tdTomato (red) in the coronal sections of LVP region of E14.5 and E16.5 in *Osr2- Cre;tdTomato* mice. Boxes in A and C are shown at higher magnification in B and D, respectively. Arrows in B and D indicate that the green MyoD or MHC signal does not overlap with the red tdTomato signal at E14.5 or E16.5. The scale bar in A indicates 500 µm in A and C; the scale bar in B indicates 100 µm in B and D.

**Figure 3—figure supplement 2.**
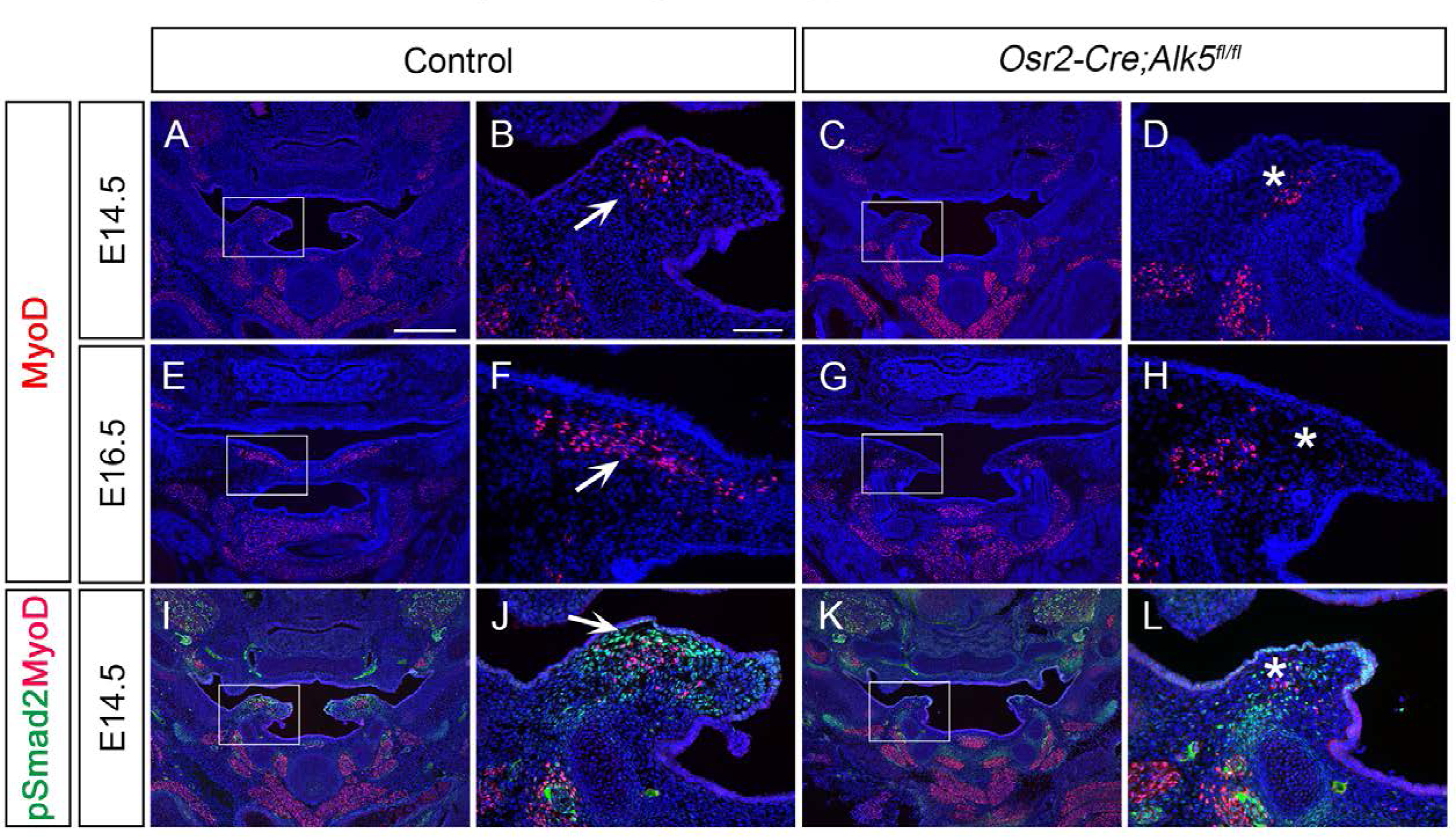
*Osr2-Cre;Alk5^fl/fl^*mice exhibit palatal shelf and myogenic defects in the LVP region from E14.5 onwards. (A-H) Immunofluorescence of myogenic marker MyoD (red) in coronal sections of palatal shelves of control mice and *Osr2-Cre;Alk5^fl/fl^* mice at E14.5 (A-D) and E16.5 (E-H) LVP region. Boxes in A, C, E, and G are shown at higher magnification in the B, D, F, and H, respectively. Arrows in B and F indicate the presence of MyoD+ cells in the control, and asterisks in D and H indicate reduced MyoD+ cells in the *Osr2- Cre;Alk5^fl/fl^* mice. (I-L) Immunofluorescence of MyoD (red) and pSmad2 (green) in the coronal section of the LVP region at E14.5. Boxed areas in I and K are enlarged in J and L, respectively. The asterisk in L corresponding to the arrow in J indicates the loss of the pSmad2 signal following disrupted TGF-β signaling. The scale bar in A indicates 500 µm in A, C, E, G, I, and K; the scale bar in B indicates 100 µm in B, D, F, H, J, and L.

**Figure 4—figure supplement 1.**
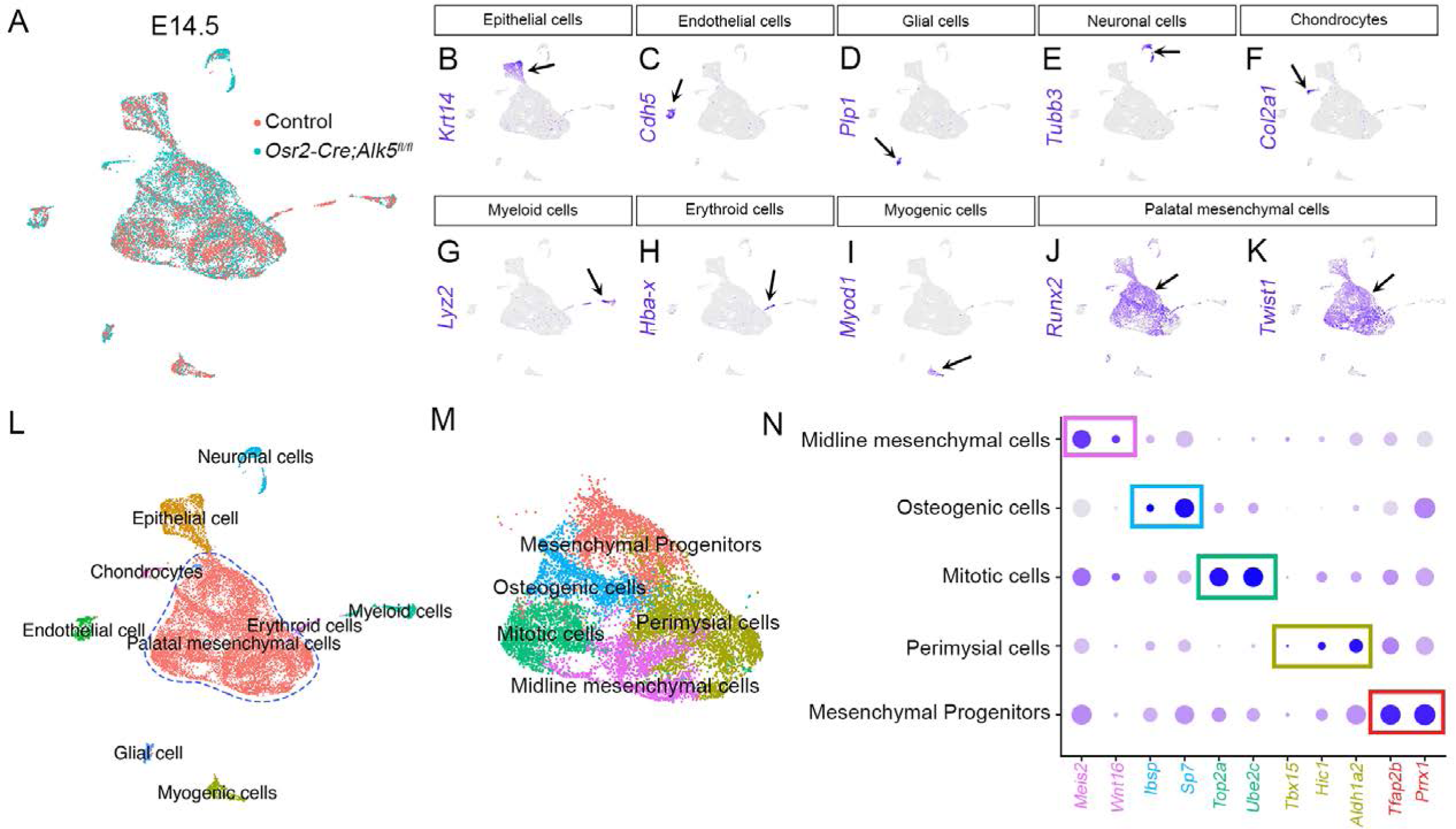
scRNAseq analysis identified distinct populations of palatal mesenchymal cells in E14.5 *Osr2-Cre;Alk5^fl/fl^* and control soft palates. (A) Integration analysis of scRNAseq from the soft palates of control and *Osr2-Cre;Alk5^fl/fl^* mice at E14.5. (B-J) Expression patterns of marker genes distinguishing non-mesenchymal cells (B-I) from palatal mesenchymal cells (*Runx2+/Twist1+*) (J-K). (L) Identification and characterization of different cellular populations in the E14.5 *Osr2-Cre;Alk5^fl/fl^* and control soft palates. (M) Individual UMAP clustering of palatal mesenchymal cells of E14.5 *Osr2-Cre;Alk5^fl/fl^* and control soft palates. (N) Marker genes from the integration of scRNAseq analysis of individual cellular clusters between *Osr2-Cre;Alk5^fl/fl^* and control mice at E14.5. Note highlighted markers for perimysial cells: *Tbx15, Hic1,* and *Aldh1a2*.

**Figure 5—figure supplement 1.**
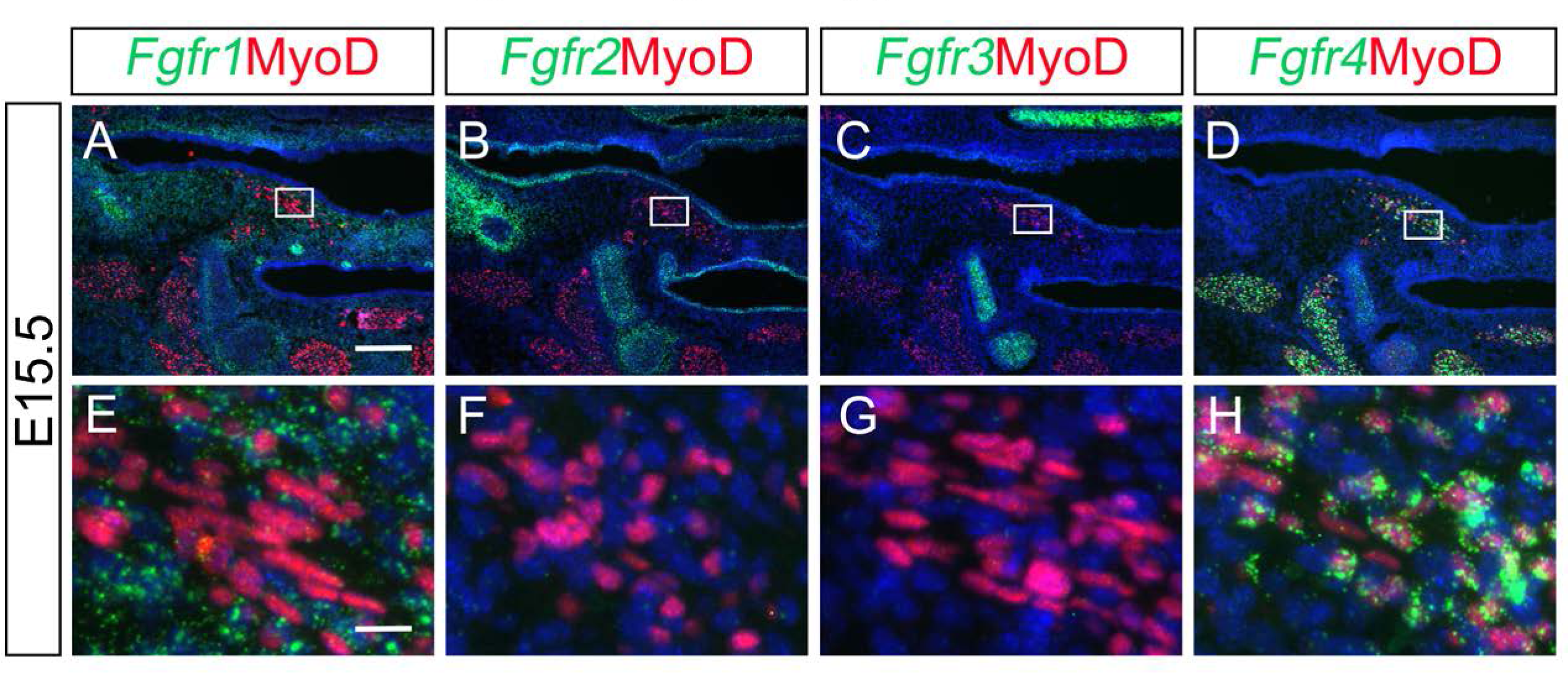
Fgf18 receptor *Fgfr4* is predominantly expressed by myogenic cells in the LVP regions. (A-H) Immunofluorescence of MyoD (red) and RNAScope *in situ* hybridization for *Fgfr1* (green) (A, E), *Fgfr2* (B, F), *Fgfr3* (C, G), and *Fgfr4* (D, H) in coronal sections of LVP region at E15.5 control mice. Boxed areas in A-D are enlarged in E-H, respectively. Scale bars in A and E indicate 200 μm for A-D and 20 μm for E-H, respectively.

**Figure 6—figure supplement 1.**
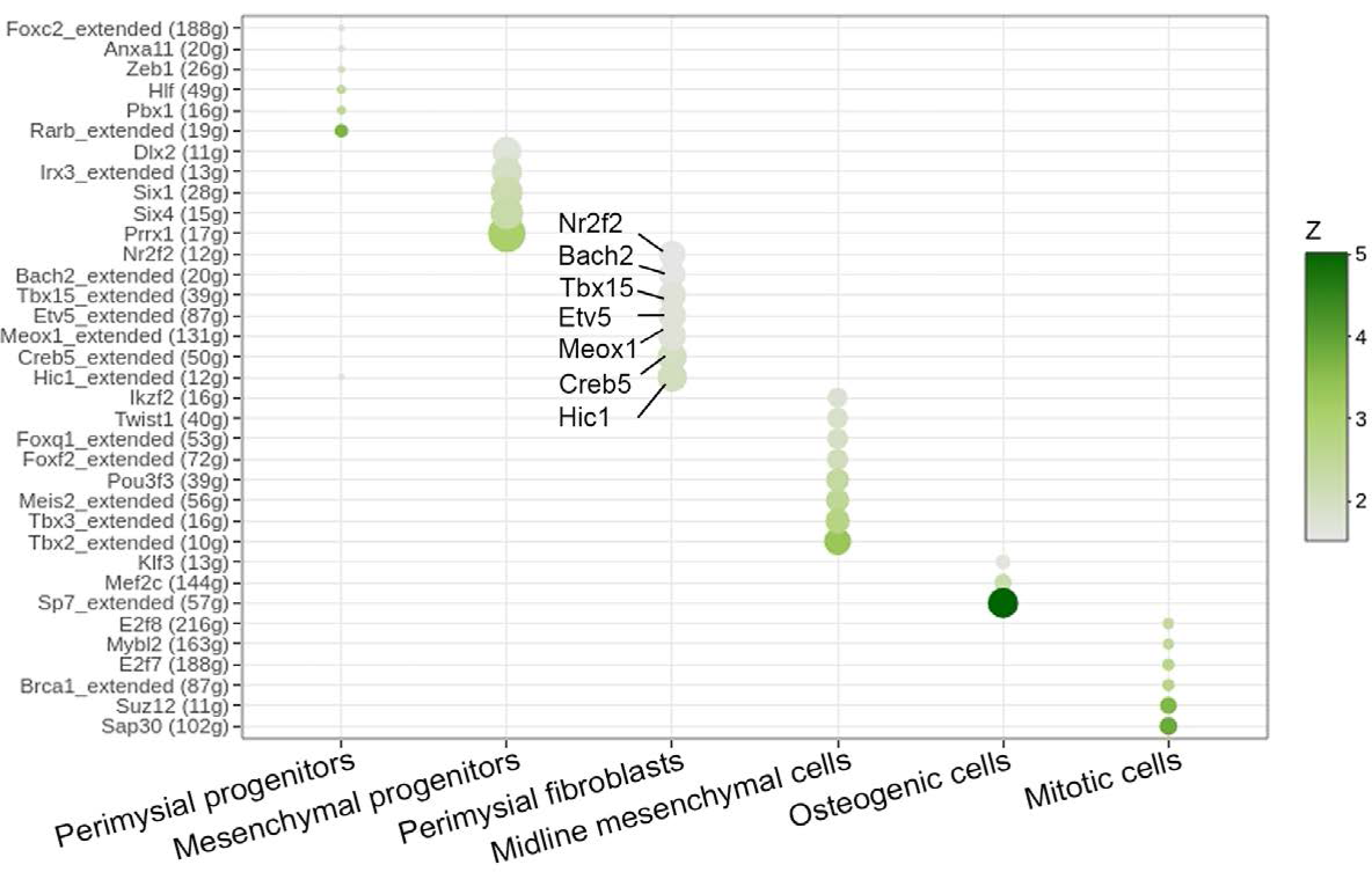
SCENIC analysis identified individual cell type-specific regulons from E13.5-E15.5 soft palatal scRNAseq data. Dot plot indicates predicated regulons for perimysial progenitors, mesenchymal progenitors, perimysial fibroblasts, midline mesenchymal cells, osteogenic cells, and mitotic cells. Note the regulons identified for perimysial fibroblasts are *Hic1*, *Creb5, Meox1*, *Etv5*, *Tbx15*, *Bach2*, and *Nr2f2*.

**Figure 6—figure supplement 2.**
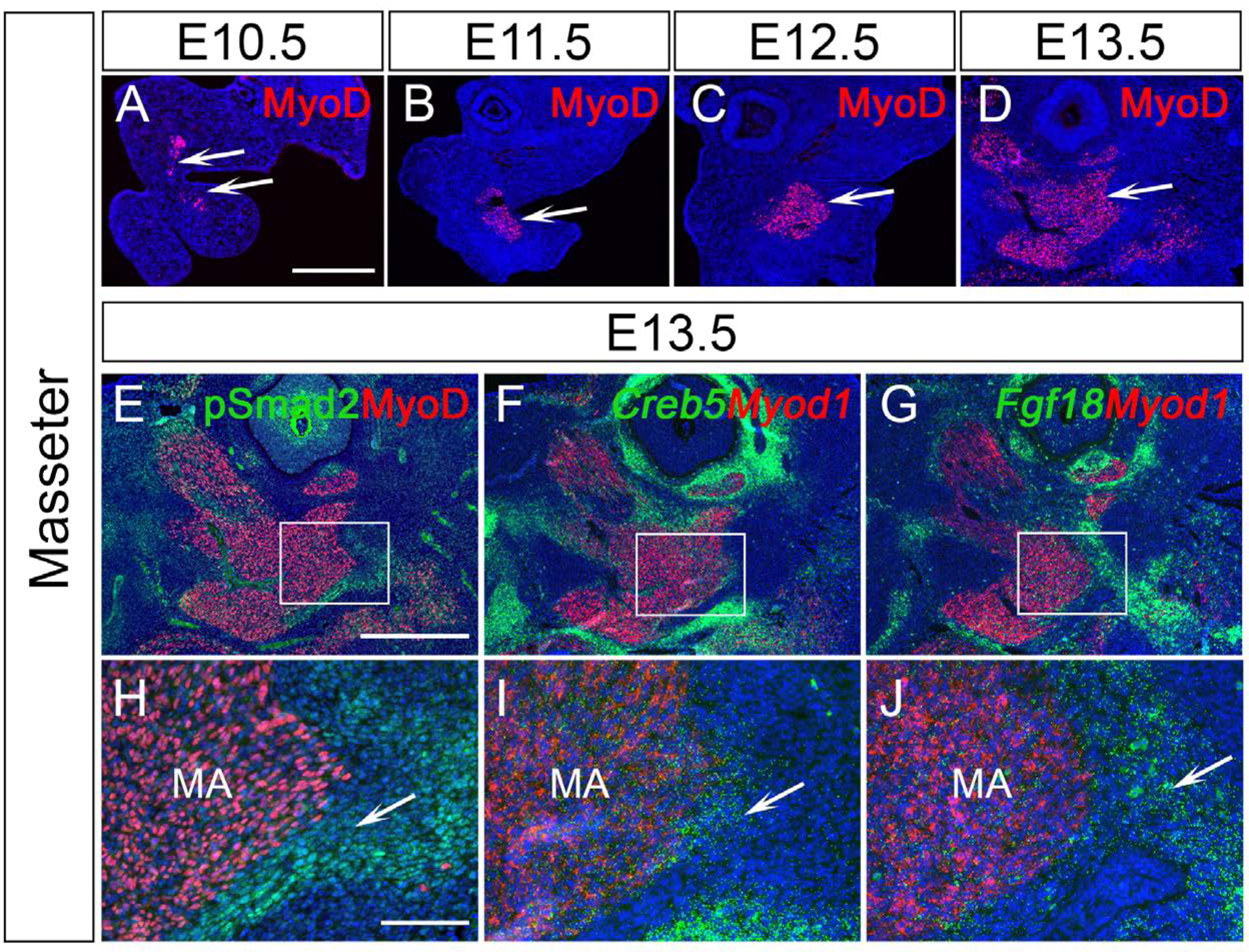
TGF-β signaling, *Creb5*, and *Fgf18* are expressed in a similar region of the perimysial cells of the masseter at E13.5. (A-D) Immunofluorescence of MyoD (red) in sagittal sections of control mouse head in the masseter region at E10.5 (A), E11.5 (B), E12.5 (C), and E13.5 (D). White arrows in A-D point to forming masseter. (E, H) Immunofluorescence of MyoD (red) and pSMAD2 (green) in sagittal sections of control mouse head in the masseter muscle region at E13.5. Boxed area in E is enlarged in H. The white arrow in H points to a positive signal. (F-G, I-J) RNAScope *in situ* hybridization for *Myod1* (red) and *Creb5* (green) (F and I) or *Fgf18* (green) (G and J) in sagittal sections of control mouse head at in the masseter region E13.5. Boxed areas in F and G are enlarged in H and I, respectively. White arrows in I and J point to positive signals. MA, masseter. Scale bars in A and E indicate 500 μm for A-D and E-F, respectively; scale bars in H indicate 100 μm in H-J.

**Figure 7—figure supplement 1.**
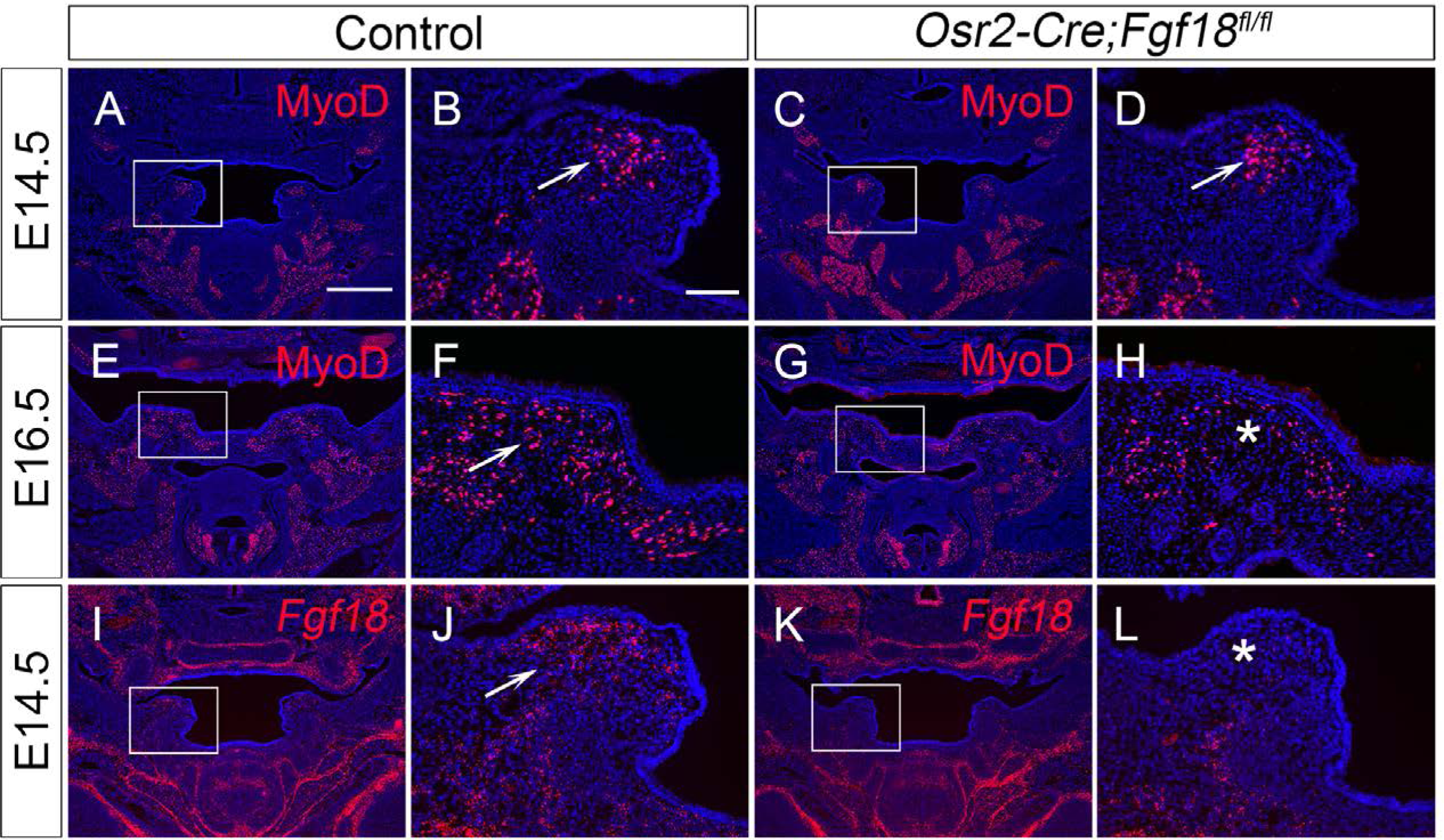
*Osr2-Cre;Fgf18^fl/fl^*mice exhibit myogenic defects during LVP development. (A-H) Immunofluorescence of MyoD (red) in coronal section of LVP region at E14.5 (A-D) and E16.5 (E-H) from control (A-B, E-F) and *Osr2-Cre;Fgf18^fl/fl^*mice (C-D, G-H). Boxed areas in A, C, E, and G are enlarged in B, D, F, and H, respectively. White arrows in B, D, and F point to the presence of MyoD+ cells. The white asterisk in H indicates a decrease of MyoD+ cells in E16.5 *Osr2-Cre;Fgf18^fl/fl^* mice. (I-L) BaseScope *in situ* hybridizations of *Fgf18* Exon1C (red) in the coronal section of LVP region at E14.5 from control (I-J) and *Osr2- Cre;Fgf18^fl/fl^* mice (K-L). Boxed areas in I and K are enlarged in J and L, respectively. The white arrow in J points to a positive signal, and the asterisk in L points to a reduced signal. Scale bar in A indicates 500 μm in A, C, E, G, I, and K; scale bar in B indicates 100 μm in B, D, F, H, J, and L.

